# Context-dependent effects of mutations on complex splicing decisions

**DOI:** 10.1101/2025.03.06.641829

**Authors:** Congxin Li, Sebastian Unic, Monika Kuban, Mirko Brüggemann, Julian König, Kathi Zarnack, Stefan Legewie

## Abstract

Alternative pre-mRNA splicing is an essential step in human gene regulation, and mutation-induced aberrant splicing is frequently found in diseases and therapy resistance. Splicing regulation is highly dependent on sequence and cellular context, posing a challenge to predict outcomes of splicing-related disease mutations. Here, we use kinetic modeling to derive the underlying quantitative principles, describing splice site competition and downstream effects on a wide spectrum of splice isoforms. Employing statistical learning on a large-scale mutagenesis dataset for *CD19* mRNA splicing, our model quantitatively describes the generation of 93 RNA isoforms. It takes into account various splicing events such as cassette exon skipping, intron retention, and extensive alternative 3’ and 5’ splice site usage, which are implicated in CART therapy resistance in leukemia. Beyond *CD19*, by analyzing genome-wide RNA sequencing data and large-scale screening of synthetic splicing decisions, we find that splice site distance is an important parameter controlling the switch from canonical to alternative/cryptic splice site usage, as mutations located in between two nearby splice sites show a pronounced directionality, regulating up- and downstream effectors in opposite direction. Taken together, our work demonstrates that a quantitative description of splice site competition provides insights into context-dependent, complex isoform changes and cryptic splice site activation in health and disease.

## Introduction

Most eukaryotic genes carry discontinuous information, consisting of (protein-)coding exons interspersed by non-coding introns. During mRNA maturation, introns are removed in a process called splicing. The modular organization of the genome allows for alternative splicing, the potential formation of multiple different mRNA isoforms from the same transcript, allowing for greater proteome diversity without the need for a higher gene number^1–3^. While alternative splicing is required for several physiological processes^4,5^, such as embryogenesis, cell differentiation and adaptive immune response, aberrant splicing is frequently involved in the development of complex diseases such as cancer and neuromuscular disorders and thus remains a target of intensive research^6,7^.

During the past decades, a large body of work identified the molecular components of the splicing machinery^8^. Splicing is catalyzed by a large macromolecular complex, the spliceosome, which assembles on *cis*-regulatory sequence elements marking the exon boundaries, the so-called 3’ and 5’ splice sites. Since splice-site sequences are highly degenerate^9,10^, additional *cis*-regulatory elements are required to enhance spliceosome assembly on genuine splice sites (exonic and intronic enhancers) or to repress non-specific spliceosome assembly (exonic and intronic silencers). These enhancers and repressors are recognized by *trans*-acting RNA binding proteins (RBPs) that in turn strengthen or weaken spliceosome recruitment via protein-RNA and proteinprotein interactions or molecular competition, respectively^11^. Alternative splicing occurs by the modulation of splice site recognition, which leads to various outcomes. Although these outcomes occur in complex combinations, they are often described in terms of binary splicing events, e.g., the inclusion vs. skipping of cassette exons, the use of canonical vs. alternative 3’ and 5’ splice sites or the splicing vs. retention of introns.

The principles of alternative splicing regulation by *cis*-regulatory elements and *trans*-acting RBPs, commonly known as the splicing code, have been characterized by various experimental and theoretical methods. For individual splicing decisions, high-throughput mutagenesis approaches determined the effect of all possible point mutations on splicing outcomes^12–15^, or explored a larger sequence space by characterizing completely degenerate sequence regions nearby splice sites^16^. Genome-wide RNA sequencing data simultaneously quantifies splicing outcomes of all exons in a cell, and when combined with knockdown approaches yields global views RBP-mediated splicing regulation^17,18^. Based on such genome-wide datasets, machine learning models have been trained to predict splicing outcomes from sequence, focusing on cassette exon inclusion/skipping^19,20^, intron retention^21^, or alternative splice site usage^22,23^. Novel frameworks in artificial intelligence promote the scalability and interpretability of these computational algorithms^24,25^.

Mutations in *cis*-regulatory sequence elements and *trans*-acting RBPs commonly perturb splicing outcomes in cancer, with implications on disease progression and therapy relapse^12,13^. Since disease phenotypes typically manifest themselves in certain tissues only, it remains a challenge to predict how mutation effects on splicing depend on cellular context, e.g., the expression levels of splice-regulatory RBPs. Mathematical models describing the kinetics of splicing reactions yielded quantitative insights into such context-dependency, explaining why *cis*- or *trans*-acting perturbations affect certain exons or exon variants more than others^12,14^. By taking into account splice site competition, these models revealed that cassette exons with intermediate inclusion levels show the strongest splicing changes, suggesting that the starting inclusion level is a quantitative predictor of mutation effects on cassette exon inclusion. Another benefit of kinetic models is their scalability to multiple isoforms. By simulating the spliceosomal recognition of multiple exons in a transcript, it is possible to derive a quantitative framework describing the synthesis of many alternative isoforms and their coordinated regulation by perturbations^26–29^. Hence, while most experimental studies and sequence-based splicing models focused on binary splicing decisions, kinetic splicing models are a promising tool to explore the full complexity of multi-dimensional splicing outcomes. However, to date, their application was limited to data with low isoform complexity, raising the question of whether kinetic splicing models can also capture highly complex datasets combining various types of splicing outcomes.

Along these lines, we derive here a complex kinetic splice site competition model simultaneously accounting for cassette exon inclusion, alternative 3’ and 5’ splice site choice and intron retention. We train this model using a complex *CD19* random mutagenesis dataset, in which the genesis of 93 splice isoforms is measured for thousands of sequence variants. We show that the model quantitatively captures the context-dependence of mutation effects on the full isoform spectrum and uses a hybrid regression approach to infer single mutation effects based on the measured mutation combinations. We then apply the model to learn the sequence determinants of alternative 3’ and 5’ splice site choice, and identify two key factors, splice site distance and directionality in splice site regulation: in both *CD19* and genome-wide datasets, alternative splice sites near the canonical one are more likely to be activated, because mutations simultaneously regulate nearby up- and downstream splice sites in opposite direction. In contrast, far-away splice sites are regulated independently, implying that switching from one splice site to another is less effective. Our model is readily applicable to other complex splicing decisions and provides a generic framework to connect the *cis*-regulatory sequence context to isoform diversity in health and disease.

## Results

### Molecular competition explains context dependence of alternative splice site choice

Human exons are frequently characterized by alternative 3’ or 5’ splice sites which can give rise to shorter and longer exon variants. Splice-site choice is generally thought to be shaped by the competition of alternative splice sites. To quantitatively understand this competition and its modulation, we analyzed a simple kinetic model in which an exon has two possible 3’ splice sites (Fig. 1a). We termed the most commonly used splice site can-3’SS (canonical) and the less used alt-3’SS (alternative).

**Fig. 1.**
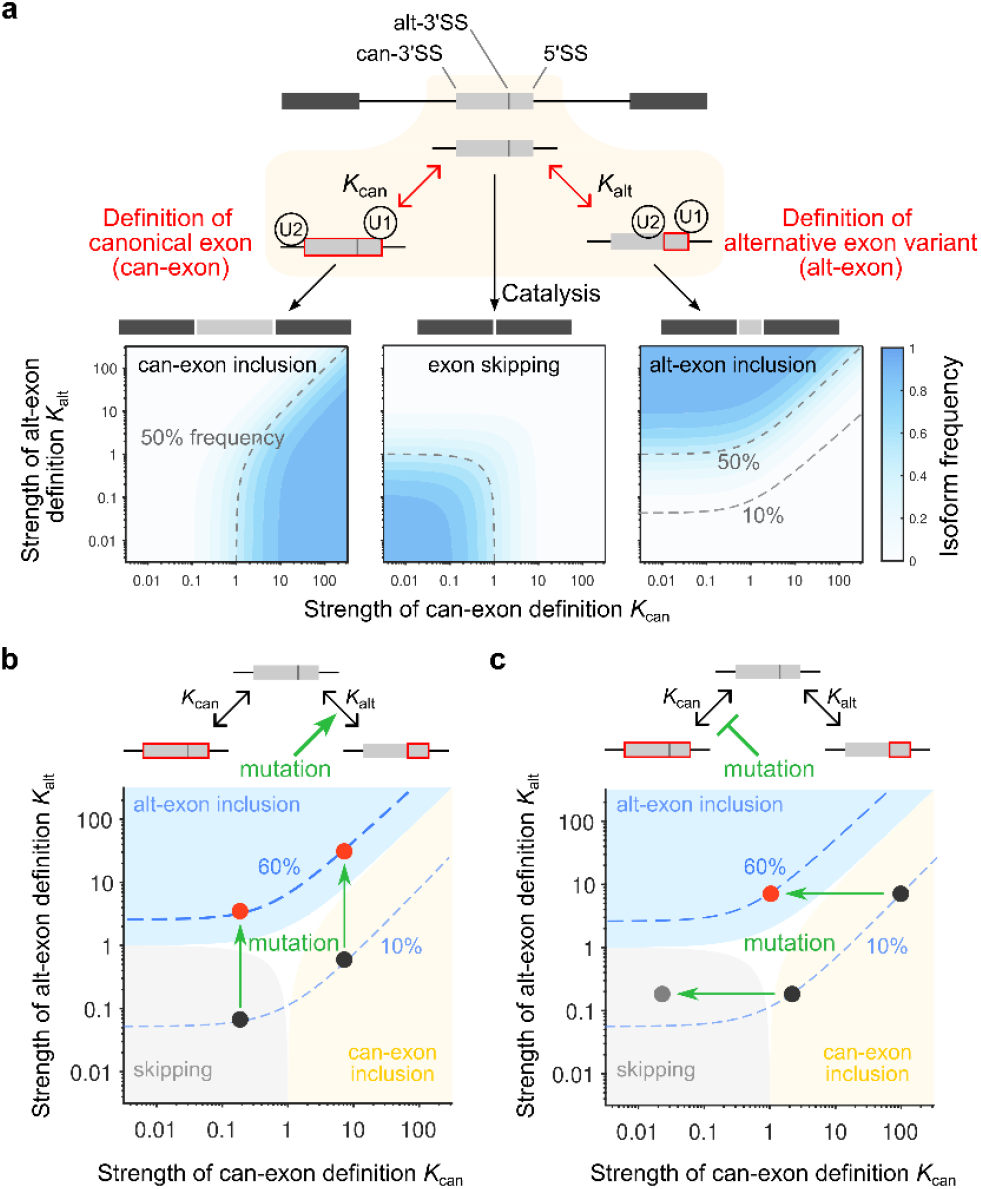
Kinetic competition model describes context-dependent alternative splice-site choice. **a** A minimal model of can-3’SS vs. alt-3’SS choice. Top: Mutually exclusive definition of canonical exon and its alternative variant (can-exon and alt-exon, respectively) by competitive and reversible binding of a cross-exon U1-U2 complex to two possible 3’SSs and a common 5’SS with the affinities *K*_can_ and *K*_alt_. In the subsequent irreversible splicing catalysis step, can-exon and alt-exon inclusion isoforms are generated, or the whole exon is skipped if undefined. Bottom: Isoform landscapes of the competition model for varying exon definition parameters *K*_can_ and *K*_alt_. Color code: isoform frequency; dashed line: 50% (and 10% for alt-exon) isoform frequency isoline. **b** and **c** Context-dependence of mutation effects on alt-exon isoform activation. Starting from an unmutated wildtype (black circle) a mutation (green arrow) can shift splicing outcomes into a new regime (red circle), and the outcomes depend on the starting isoform distribution (‘context’) and the mutation strength (length of arrow). (**b**) The arrows show the effect of a strong mutation which increases alt-exon recognition (*K*_alt_) to activate the alt-exon inclusion isoform (e.g., from a frequency of 10% to 60%; blue dashed lines) for varying initial isoform distributions. (**c**) Strong mutations diminishing the recognition of the competing canonical exon (*K*_can_) lead to either skipping or alt-3’SS activation starting from the same alt-exon isoform frequency (10%), depending on the chosen alt-exon recognition strength *K*_alt_. Blue dashed line: 10% or 60% frequency isoline for alt-exon isoform. Shades: parameter areas giving rise to high frequencies (≥ 50%) for skipping (grey), can-exon (yellow) and alt-exon (blue) inclusion isoforms.

Splicing in mammalian cells predominantly occurs via an exon definition mechanism, in which the pioneering subunits of the spliceosome (U1 and U2) initially form cooperative complexes across exons^30^. We therefore assumed that the exon is defined as a functional unit as in our previous modeling work^28^. During exon definition, we considered that the can-3’SS and alt-3’SS compete for pairing with a shared 5’ splice site (Fig. 1a). In terms of splicing outcomes, such competition leads to the recognition (definition) of either the canonical exon (termed can-exon) or its alternative variant (termed alt-exon); or the exon is skipped if neither exon variant is defined, resulting in three possible splicing outcomes. Our model can readily be extended to describe more complex splicing decisions, e.g., by considering alternative 5’ splice site usage (Methods), as discussed in later sections.

To quantify the competition between the two exon variants, we introduced two model parameters *K*_can_ and *K*_alt_, which reflect the affinity of early spliceosomal U1-U2 complex binding to the can-exon and alt-exon, respectively. The binding affinities reflect the probability of spliceosome assembly and they may be modulated by various molecular determinants, such as splice-site strength, sequence context and mutations. To gain insights into splicing regulation, we derived an abundance landscape for each mRNA isoform at steady state as a function of the two parameters (Fig. 1a, lower panels; Methods). Depending on parameter choice, splicing is dominated by exon skipping (both *K*_can_ and *K*_alt_ low), canonical exon inclusion (*K*_can_ >> *K*_alt_) or alternative exon inclusion (*K*_can_ << *K*_alt_) (Fig. 1a, lower panels).

We next explored ways how sequence mutations can promote the use of the alt-3’SS. An intuitive regulatory mode is to strengthen the binding affinity of the alt-exon variant, i.e., increasing *K*_alt_ (vertical, up-pointing arrows in Fig. 1b). In principle, such mutations can always increase the level of the alt-exon variant, irrespective of the starting point in the parameter landscape, provided the fold-change of *K*_alt_ is large enough. However, depending on the starting point, the fold-change of *K*_alt_, reflected by the length of the mutation arrow in Fig. 1b, needs to be adjusted to achieve a certain level of alt-exon inclusion (Fig. 1b; Supplementary Fig. 1a and b).

This starting point dependency of mutation effects is consistent with previous work on cassette exon inclusion, where the starting percent spliced-in index (PSI = inclusion/(inclusion + skipping)) was shown to determine the PSI change elicited by a mutation^12,14^. However, in our three-dimensional splicing landscape, the same shift from a low (10%) to a high (60%) alt-exon inclusion frequency can be observed for different starting levels of can-exon inclusion and exon skipping levels. Furthermore, depending on these starting points in the 2D landscape, the mutation induces distinct changes in can-exon inclusion and exon skipping (Figs. 1a and b). This suggests that a single metric (PSI or a particular isoform frequency) does not capture the full complexity of splicing decisions with more than two outcomes. Therefore, multi-dimensional splicing landscapes are essential to quantitatively interpret mutation effects on alternative splice-site usage.

Another mode of activation for the alt-exon is to weaken the competing can-exon by mutation (lowering *K*_can_; horizontal, left-pointing arrows in Fig. 1c). Interestingly, this modulation is only effective when both starting exon definition parameters (*K*_can_ and *K*_alt_) are sufficiently high, i.e., if can-exon and alt-exon are efficiently recognized in the absence of mutation. In contrast, the same mutation exclusively induces skipping if the starting *K*_can_ and *K*_alt_ are both low. Furthermore, both alt-exon (upper arrow) and skipping accumulation (lower arrow) by a given mutation can happen at the same starting frequency of the alt-exon (e.g., 10% in Fig. 1c). Thus, the starting frequency of the alt-exon isoform no longer predicts its accumulation upon mutation and quantitative knowledge about the recognition strength of both exon variants is necessary to predict the splicing outcomes (Methods; Supplementary Fig. 1). This further corroborates that a single metric like the PSI cannot capture the complexity of splicing outcomes when more than two splice isoforms are produced.

Taken together, these simulations suggest that kinetic models simultaneously taking into account multiple splice isoforms are required to fully comprehend the quantitative principles of alternative splice-site choice and its modulation by sequence mutations.

### Splice-site competition quantitatively describes large-scale *CD19* mutagenesis data

To confront the splice-site competition theory with data, we made use of our recent work where we comprehensively characterized splicing decisions of a *CD19* minigene reporter comprising exons 1-3 in the NALM6 B-cell line (Fig. 2a; Supplementary Fig. 2). Using random mutagenesis, we established over 9600 minigene variants, each harboring 9.7 point mutations on average, which were transfected into cells and the splicing outcomes were quantified by RNA sequencing^13^. To robustly quantify the splicing outcomes, we filtered out the minigenes with low sequencing read count (< 100) and keep ∼ 9000 minigene variants (8996 in replicate 1 and 8597 in replicate 2) for further study. The large-scale screening revealed an extensive usage of alternative/cryptic 3’ and 5’ splice sites (Supplementary Fig. 2a). In the following, we will use the term alternative for those splice sites used in the unmutated wildtype *CD19* minigene, while the notation cryptic refers to splice sites emerging only in mutant *CD19* minigenes. *CD19* exon 1 is characterized by one canonical and six cryptic 5’ splice sites, resulting in a total of seven exon 1 variants. Exon 2 has six possible 3’ splice sites (1 canonical, 1 alternative and 4 cryptic) and 24 5’ splice sites (1 canonical and 23 cryptic), which may combine into 144 (6×24) exon variants. However, only 34 variants were observed for exon 2, indicating weak paring between the majority of 3’ and 5’ splice sites. In exon 3, 34 3’ splice sites (1 canonical, 1 alternative and 32 cryptic) lead to 34 exon variants.

**Fig. 2.**
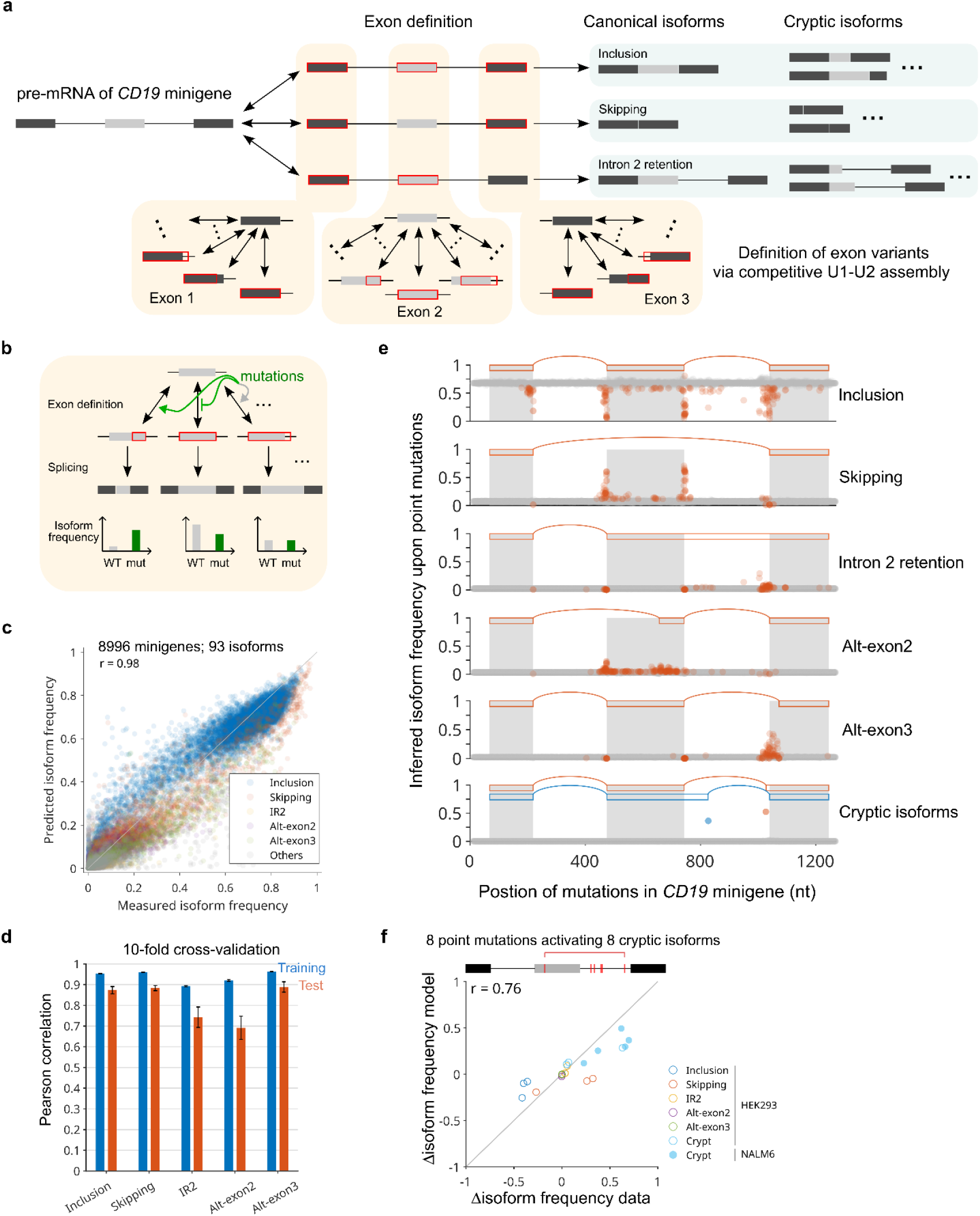
Multi-dimensional splice-site competition model quantitatively describes largescale *CD19* mutagenesis data. **a** Multi-dimensional splice-site competition model describes the generation of 93 splice isoforms in the *CD19* minigene. In the model, the three minigene exons are defined independently by reversible U1-U2 binding (indicated by red boxes). Shown are three exemplary exon definition states, which give rise to exon 2 inclusion (all exons defined), exon 2 skipping (exon 2 undefined) and second intron retention (exon 3 undefined) as splicing outcomes. As an additional layer of complexity, multiple competing exon variants, corresponding to alternative 3’SS or 5’SS, show mutually exclusive definition, leading to the generation of alternative/cryptic isoform variants. Based on the *CD19* data, we considered 7, 34 and 34 variants (including canonical exon) for exon 1, 2 and 3, respectively (see Supplement for details). **b** Inference of single point mutation effects from large-scale mutagenesis data by fitting the simulated splice isoform frequencies to the measured ones under the assumption that mutations affect the exon variant definition rates. Since multiple mutations are assumed to act additively, a linear regression approach was formulated to infer single point mutation effects on exon definition from measurements of minigenes harboring multiple mutations (see Methods for details). **c** Linear regression approach faithfully captures the frequencies of 93 isoforms in ∼ 9000 (8996 in replicate 1 and 8597 in replicate 2) randomly mutated *CD19* minigenes (Pearson correlation *r* = 0.98). The isoform frequencies fitted by the model are plotted against the measured datapoints. Each dot represents one minigene variant and the colors reflect splice isoform identity. **d** The predictive power of the model was validated by 10-fold cross validation, in which the set of ∼ 9000 minigene variants (8996 in replicate 1 and 8597 in replicate 2) was split ten times into 90% of the minigene variants used for fitting (‘training’) and remaining 10% used for validation (‘test’). Bars show the mean Pearson correlation coefficient as measure of model-data coherence across the 10 data partitions, and the error bars reflect its standard deviation. **e** Model-inferred frequencies of the five major (1st – 5th rows) and two representative (bottom row) cryptic *CD19* minigene splice isoforms for hypothetical single-mutation minigenes. Shown is the inferred isoform frequency for each single mutation (red dots), sorted according to mutation position in the *CD19* minigene. For reference, the canonical exons are indicated by grey shading, the red rectangles and splicing arcs represent the exons used in the considered isoform. **f** Experimental validation of predicted single-mutation effects on cryptic isoforms. 8 targeted single point mutations were introduced into the *CD19* minigene in NALM6 or HEK293 cells, and isoform frequency (IF) changes (ΔIF = IF_mut_ - IF_wt_) predicted by the model are compared to splicing measurements by capillary electrophoresis. The previously published NALM6 data^13^ covers only the predicted accumulating cryptic isoform, while the Hek 293 data additionally covers all five major *CD19* splice isoforms. *r*: Pearson correlation coefficient.

The combination of these 75 variants of the three CD19 exons can, in theory, generate more than 9000 splice isoforms (see Method). Nonetheless, only 93 isoforms were detected in the actual splicing measurements of the *CD19* minigene library (Supplementary Fig. 2b). Five isoforms were already found for the unmutated wildtype minigene^13^. Three of them use only canonical splice sites, i.e., exon 2 inclusion, exon 2 skipping and second intron retention. Two are inclusion isoform variants, respectively using alternative 3’ splice sites in exon 2 (at position 658) and exon 3 (at position 1073), termed alt-exon2 and alt-exon3. The remaining 88 isoforms are termed cryptic isoforms below, as they were absent in the wildtype, but accumulated in at least one mutated minigene with maximal frequencies ranging from 5% to 90%^13^. Given this high number of alternative splicing events, the *CD19* dataset is well suited to test the splice site competition model (Fig. 1).

To describe the generation of such a variety of isoforms, we embedded the splice-site competition model (Fig. 1) into a previously derived model that describes cassette exon splicing in a three-exon minigene^28^ (Fig. 2a). Within each exon, multiple variants compete for U1-U2 binding and thus they are recognized (or defined) in a mutually exclusive manner. The probability of a certain exon variant being defined within an exon is determined by the relative affinity of that variant to the U1-U2 complex. For example, the recognition probability of the *i-*th variant of a given exon *p*_*i*_ = *K*_*i*_ / (1 + *K*_1_ + *K*_2_ + … + *K*_n_) with *i* = 1, …, *n*, where *K*_i_ is the binding affinity of the *i*-th variant and *n* is the total number of variants for that exon (derivation see Methods). The sum of these recognition probabilities (∑*p*_i_) determines whether the exon is used at all.

To link spliceosome binding to splicing fates, we assumed that the set of defined exons determines splicing outcomes: For instance, the binding states 111 (binary notation for all three exons recognized), 101 (middle exon not recognized) and 110 (last exon not recognized) give rise to exon 2 inclusion, skipping and second intron retention isoforms, respectively (Fig. 2a). For each of these fates, multiple exon variants were considered, i.e., 7, 34 and 34 variants of exon 1, 2 and 3 are combined for inclusion (111) with recognition probabilities *α*_i_, *β*_i_ and *γ*_i_ (defined as above for *p*_i_). For example, the recognition probability of the *i*-th variant of exon 1 is *α*_*i*_ = *K*_1*i*_ / (1 + *K*_11_ + *K*_12_ + … + *K*_17_), where *K*_1*i*_ is the affinity of spliceosome binding to this exon variant. Together with splicing fates other than inclusion (skipping, first intron retention, second intron retention, full intron retention), our model can give rise to a total pool of 9189 isoforms (Methods).

To learn the parameters (*K*_i_) for exon variant recognition from isoform measurements, we devised a model fitting approach based on maximum likelihood estimation by which we could analytically calculate the parameters based on the sequencing read counts of 93 isoforms in ∼ 9000 (8996 in replicate 1 and 8597 in replicate 2) minigenes (see Methods). Based on these parameter estimates, we performed forward simulations to isoform abundances to compare the modeling result to the original experimental data used for fitting (Supplementary Fig. 3). We observed a strong correlation (*r* = 0.95) between model and data across all ∼ 800,000 datapoints (*n* = 836,628 in replicate 1 and *n* = 799,521 in replicate 2). This suggests that a model with only two elementary rules, i.e., independent definition of exons 1-3 and within-exon competition of variants, quantitatively describes the *CD19* high-throughput experimental data.

Remarkably, the model explains the small number of actually realized isoforms in the data (93 isoforms) compared to the total possible number of isoforms (*n* = 9189). This discrepancy solely arises due to certain exon variants being weakly recognized and therefore extremely unlikely to occur in combination in the experimentally observed isoform space.

### Model predicts single and combined mutation effects on splice-site choice

Having determined the relative strength of exon variants for combined mutations (minigenes), we sought to infer the impact of individual point mutations on exon variant recognition. To do so, we adopted our previously described linear regression approach, in which we describe the combined mutation effect as the sum of underlying individual point mutations^12,13^.

In the context of the splice-site competition model, we assumed additivity of mutations on spliceosomal binding energies (Fig. 2b and Methods). Specifically, we assumed that the unknown log-fold-effects of individual mutations on the exon variant recognition parameters (*K*_i_) sum up to the total log-fold-change of combined mutations in each minigene (see Methods). To implement this rule, we derived a set of ∼ 9000 equations (8996 in replicate 1 and 8597 in replicate 2), one for each minigene, which related the measured combined mutation effect to the unknown sum of individual mutation effects. By fitting this set of equations to the experimental data (Fig. 2c), we inferred the unknown individual mutation effects on the spliceosome binding parameters and thus on splice isoform frequencies. Since many of the point mutations likely have no effect on splicing, we employed a penalized linear regression approach and optimized the predictive power of the model using 10-fold cross-validation (see Methods). The final regression model provided an excellent fit to the training data (*r* = 0.89 - 0.96 across the 93 isoforms) and accurately predicted unseen test data (*r* = 0.69 - 0.89) (Fig. 2d). This confirmed that mutations show log-additive behavior at the level of exon definition.

The regression model predicted single mutation effects on the five major *CD19* isoforms, which is in line with our previous work^13^, and it further identified potential point mutations that strongly activate rare cryptic isoforms (Fig. 2e; Supplementary Fig. 4). To test the model-predicted activation of alternative/cryptic splice sites experimentally, we performed targeted mutagenesis introducing single point mutations into *CD19* exons 2 and 3, thereby obtaining novel 47 minigene variants (Supplementary Table 1). These single-mutation minigenes were transfected either into NALM6 cells, in which the mutagenesis screen had been performed (published data by Cortés-López et al.^13^, 5 mutants), or into HEK293 cells (this publication, 42 mutants), and analyzed for splicing outcomes using capillary electrophoresis (Supplemental Fig. 5). For 39 single mutants in HEK293 cells, the observed changes in isoform frequency agreed well with model predictions for the five major isoforms (*r* = 0.78; Supplementary Fig. 5e). Furthermore, 8 tested point mutations (5 in NALM6 and 3 in HEK293 cells) indeed led to the activation of the predicted cryptic isoforms, and the abundance of these was in quantitative agreement with the model (*r* = 0.76; Fig. 2f). Hence, the regression analysis faithfully captured how individual mutations modulate the recognition of exon variants, thereby providing a quantitative description of alternative splice-site choice.

### Efficient alt-exon activation by opposing mutation effects on competing splice sites

We next asked whether the model-predicted single mutation effects on spliceosome binding and splicing outcomes were in line with the biological expectations. To test this, we related the location of point mutations within the *CD19* exon 2 to their predicted splicing effects in NALM6 cells, projecting the model-predicted effect of each mutation on the two-dimensional splice-site competition landscape (Fig. 3a, top panels). Thereby, we focused our analysis on the usage of canonical and alternative exon 2 splice sites, while neglecting effects on other *CD19* splice sites. As expected, mutations nearby the upstream canonical splice site of exon 2 reduced the recognition of the canonical exon 2 (*K*_can_), mostly leading to skipping (top left panel). In contrast, mutations nearby the alt-3’SS at position 658 led to improved recognition of the alternative exon 2 variant and to moderate accumulation of the corresponding splice isoform (alt-exon2 isoform, two top middle panels). Moreover, the 5’ splice site of exon 2 which is shared by the canonical and alt-exon2 isoforms lowered the predicted spliceosomal binding to both exon variants as expected, again shifting the system towards exon 2 skipping (top right panel). Hence, our model maps mutations to the binding landscape of exon variants in the expected way.

**Fig. 3.**
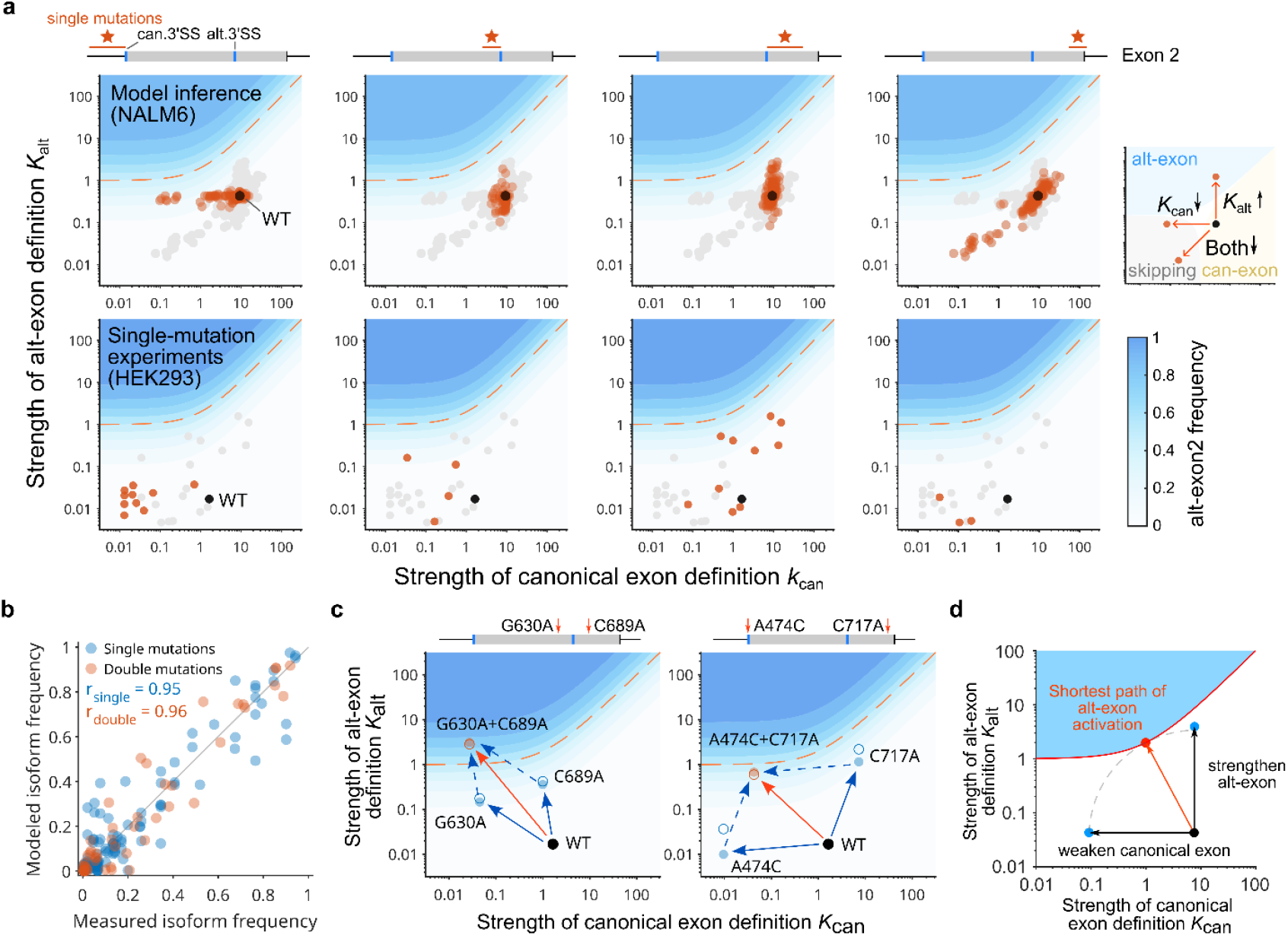
Conserved position-dependence and additivity of mutation effects across cell types. **a** Position-dependent effects of point mutations on the recognition of canonical and alternative versions of *CD19* exon 2 (alt: alt-3’SS at position 658). Top row: model-inferred exon definition rates *K*_can_ and *K*_alt_ for hypothetical single-mutation minigenes, derived by fitting the high-throughput mutagenesis data in NALM-6 cells and plotted into the 2D landscape showing the altexon2 isoform frequency (compare Fig. 1; see Methods for details). Bottom row: experimentally measured splicing outcomes of single mutation minigenes in HEK293 cells, projected on the same 2D landscape. Grey circles: all predicted/tested single mutations. Red circles: single mutations in specific regions indicated by the red horizontal bar in the exon scheme on top of each panel. **b** Single-mutation effects on can-exon and alt-exon recognition of *CD19* exon 2 are additive in HEK293 cells. Experimental splicing data (Fig. 2e) for *CD19* minigenes harboring single mutations or their combinations (15 pairs) were fitted jointly using a reduced linear regression approach covering the 5 major splice isoforms (Fig. 2; Supplementary Fig. 6). The model simultaneously captures both single- and double-mutation data, thereby suggesting additivity. See Supplementary Fig. 6 and Supplementary Table 1 and 2 for data summary. **c** Examples of additive mutation effects in HEK293 cells, shown in the 2D landscape quantifying can-exon2 and alt-exon2 definition. In both cases, one mutation mainly strengthens the alt-3’SS (blue circles mainly shifting upward; C689A in the left panel or C717A in the right panel) compared to the unmutated WT (black circle). The other mutation weakens the competing canonical splice site (blue circles mainly shifting to the left; G630A in the left panel or A474C in the right panel), resulting in the efficient activation of the alt-3’SS when the two mutations are combined (orange circles). Model fit (panel b): filled circles and arrows; Experimental data: open circles. **d** Model predicts optimal antagonistic regulation of can-3’SS and alt-3’SS for efficient activation of alt-exon2. Starting from the unmutated WT (black circle), a unique shortest path exists (red arrow) to achieve a certain alt-3’SS usage (here: 50%, red line), which combines alt-3’SS up- and canonical site downregulation.

To further confirm these model predictions, we studied splicing outcomes in HEK293 cells, mapping 27 single point mutants in exon 2 (Supplementary Fig. 5) onto the 2D abundance landscape of alt-exon2 activation. In qualitative agreement with the NALM6 model predictions (Fig. 3a, top panels), we observed the expected relations between mutation location and splicing outcomes in these HEK293 validation experiments (Fig. 3a, bottom panels; see further below for a quantitative comparison of HEK293 and NALM6 results).

Notably, even though the single mutation effects showed the expected direction of change in the exon variant space, none of them was able to effectively activate the alt-3’SS (at position 658) leading to alt-exon2, neither in the model (Fig. 3a, top panels) nor in the experimental data (Fig. 3a, bottom panels). However, the model predicted that certain combinations of these mutations will be effective, especially those combining the previously introduced basic modes of alt-exon regulation (Fig. 1b and c), i.e., weakening the canonical exon and strengthening the alt-exon.

To test whether combined mutations can indeed more effectively generate alt-exon2, we used targeted mutagenesis, generating minigenes containing double mutations near the can-3’SS and alt-3’SS (Supplementary Table 2; Methods). As expected, such double mutations enabled strong accumulation of the alt-exon2 isoform in HEK293 cells: For instance, C717A, a mutation that strengthens the recognition of the alt-exon2 variant according to our model, effectively triggered usage of the alt-3’SS when combined with A474C that weakens the canonical exon, whereas each mutation alone was ineffective (Fig. 3c, left panel). Likewise, G630A in combination with A474C also results in strong alt-exon2 activation (Fig. 3c, right panel).

We next asked whether the observed alt-exon2 activation by combined but not single mutations can be quantitatively explained by the splice-site competition model. As described for cassette exon inclusion^12,14^, the model predicts weak mutation effects for low starting alt-exon levels, and more pronounced effects of the same mutation if another mutation had primed the minigene by promoting alt-exon usage. In the splicing landscape, such priming occurs whenever the minigene is shifted close to the blue area marking predominant alt-exon usage, making it easier for a mutation to pass the threshold for 50% isoform accumulation (orange dashed line in Fig. 3c). The regression model trained on the NALM6 mutagenesis data qualitatively agreed with the single mutation validation experiments in HEK293 cells (Fig. 3a), but the magnitude of mutation effects quantitatively differed between cell types (see below). To overcome the potential confounding factor of cell-type difference in quantitative analysis of the HEK293 data, we reduced the full exon competition model (Fig. 2a) by only considering two exon variants (can-exon and alt-exon) in exon 2. We then fit this model jointly to the abundance of the *CD19* isoforms for each pair of double mutations and the corresponding single mutations assuming the log-additivity of mutation effects at the level of exon definition (see Methods). For all 15 pairs of double mutations, the model simultaneously captured both double (*r* = 0.96) and single (*r* = 0.95) mutation data (Fig. 3b, individual examples of model fit vs. data are shown in Fig. 3c). This confirms the additivity of combined mutation effects at the level of exon definition and that splice-site competition quantitatively explains the apparent cooperativity of weak mutations.

Taken together, we found that opposing regulation of the two competing exon variants, i.e., strengthening the alt-exon2 variant and weakening the canonical exon, facilitates alt-exon2 isoform activation. In theory, such opposite regulation can also be achieved by a single mutation and our model predicts a unique shortest path by which alt-exon2 activation can be achieved by minimal free energy cost in the splicing landscape (Fig. 3d; Methods). In the following, we show that the alternative variant of *CD19* exon 3 can be indeed activated by single mutations following this optimal path.

### Exon activation potential determined by splice site distance

Our high-throughput mutagenesis in NALM6 cells shows that alt-3’SS usage (at position 1073) of exon 3 can be more readily enhanced by single mutations (Fig. 2e). Thus, *CD19* exon 3 has a higher potential to be activated by mutations compared to exon 2. To test whether this could be explained by the above theory in which a single mutation jointly modulates both the canonical and the alternative splice sites (Fig. 3d), we mapped the inferred single mutation effects in exon 3 to the splice-site competition landscape (Fig. 4a). In line with our theory, the most effective single mutations simultaneously strengthened the alt-3’SS and weakened the canonical site (middle panels in Fig. 4a), aligning closely with the optimal predicted modulation (black arrow). Similar, close-to-optimal regulation was observed for single point mutations introduced into exon 3 (Supplementary Fig. 5f). Interestingly, the most effective mutations in exon 3 were located in between the two competing 3’ splice sites, in both high-throughput and targeted mutagenesis data (middle panels in Fig. 4a and Supplementary Fig. 5f). In contrast, mutations located up- or downstream of both splice sites either regulated only the nearest splice site or regulated both splice sites jointly in the same direction (outer panels in Fig. 4a and Supplementary Fig. 5). Thus, optimal cryptic exon activation may arise because mutations control up- and downstream splice sites in opposite directions (see also below).

**Fig. 4.**
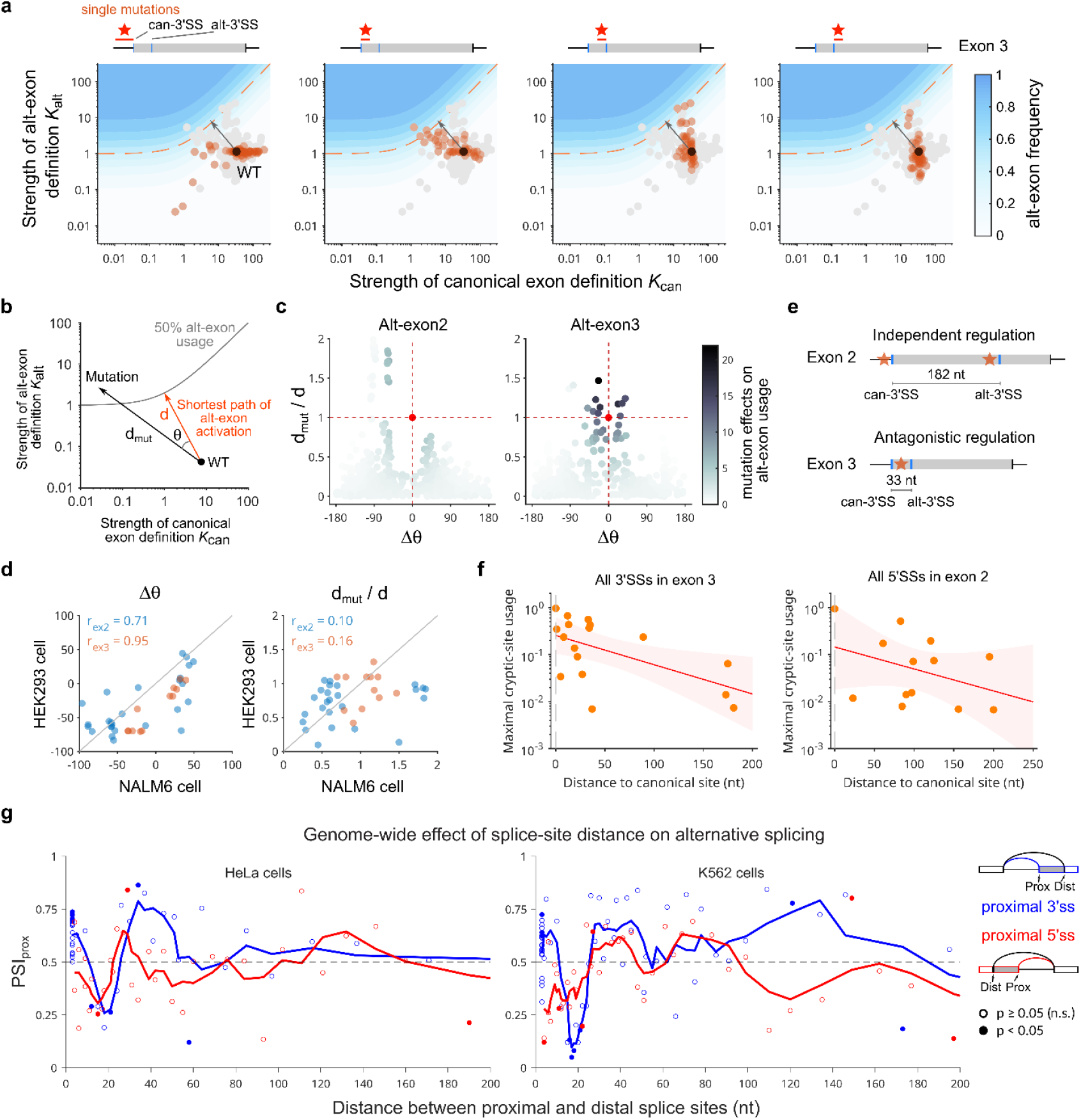
Distance from canonical site determines alternative/cryptic splice site usage. **a** Alternative splicing of *CD19* exon 3 efficiently activated by single mutations located in between the can-3’SS and alt-3’SS (alt: alternative 3’SS at position 1073). Shown are model-inferred exon definition rates (*K*_can_, *K*_alt_) for single-mutation minigenes estimated by fitting the high-throughput mutagenesis data in NALM-6 cells. Same representation as in Fig. 2a. Mutations located to blueshaded area lead to strong alt-exon 3 activation. Black arrows show the shortest path to alt-exon activation. **b - d** Short distance between splice sites in *CD19* exon 3 may allow for optimal antagonistic regulation and efficient activation of alt-exon by single point mutations (**b**) Quantification of mutation optimality in the 2D exon definition landscape using two parameters: the angle between the point mutation vector and theoretically optimal shortest path (*θ*) and the length ratio of these two vectors (*d*_mut_/*d*). (**c**) Alt-3’SS activation in exon 3 (right) shows more optimal mutation directionality (*θ*) when compared to exon 2 (left), and this leads to strong alt-3’SS usage (dark grey shading), even though mutations in both exons reach a similar maximal strength (*d*_mut_ / *d*). Same data as in Fig. 3a (top) and Fig. 4a (**d**) Short distance between can-3’SS and alt-3’SS in exon 3 (33 nt) may establish antagonistic regulation by in-between mutations, while distant splice sites in exon 2 (182 nt) are regulated independently. **e** Directionality of single-mutation effects in the exon definition landscape is consistent across cell types. Scatter plots relate mutation directionality *θ* and effect size (*d*_*mut*_*/d*) of 38 overlapping single mutations in NALM6 cells (inferred) and HEK293 cells (measured). Blue and grey data points show exon 2 and 3 mutations, respectively, and the Pearson correlation coefficients (*r*) quantify coherence of mutation effects across cell types. **f** Splice site distance globally predicts cryptic splice site activation by single mutations in the *CD19* dataset. Maximum usage of 17 alternative 3’SSs in exon 3 (left) and 13 alternative 5’SSs in exon 2 (right) across all 3688 single point mutations (inferred from NALM-6 mutagenesis) as a function of distance to the canonical splice site. Here, only splice sites harboring a consensus motif in the wt *CD19* sequence were considered (AG for 3’SS and GU for 5’SS). Straight lines and shades are fit and with confidence bands of a linear model, respectively. **g** Splice site distance has a regulatory impact on alternative 3’ and 5’ splice site choice in genome-wide RNA sequencing data from HeLa and K562 cells. Splicing events (LSVs) with alternative 3’ or 5’ splice site choice were quantified using MAJIQ^31,32^, the splice site closer to the reference exon being defined as the proximal SS (see schematic on the right). Splice site distance was quantified as the distance between proximal and distal splice site in nucleotides (nt). For each pair of proximal and distal splice sites, their relative usage was quantified as PSI_prox_ = (proximal reads) / (proximal reads + distal reads). Only splice site pairs with a comparable match to the consensus sequence, i.e. a similar MaxEnt-score^33^, were considered, requiring that |ΔMaxEnt|<2.5 (see text and Methods for details). PSI_prox_ values of alt3’ss (blue) and alt5’ss (red) events were sorted by splice site distance and binned (bin size = 20 data points), and the median PSI and mean splice site distance is plotted per bin (circles). Statistical significance of differential average proximal and distal splice site usage per bin was assessed by the Wilcoxon signed-rank test, with filled circles (p < 0.05), indicating a strong preference of one type of splice site. Due to symmetry of proximal and distal PSIs, only proximal splice site utilization is shown. A moving mean (window size = 4) was also added to improve visualization of data distribution (blue and red solid lines).

To systematically quantify the alignment with the optimal regulation vector, we calculated an angle *θ* between the observed and the optimal vector for each mutation using the NALM6 high-throughput mutagenesis model, and additionally determined the length ratio *d*_mut_/*d* between the observed and optimal vectors (Fig. 4b). In fact, for exon 3 the angle *θ* is close to zero for a subset of mutations, whereas this region tends to be excluded for exon 2, implying that the mutation-induced changes of can-3’SS and alt-3’SS recognition are close to optimal in exon 3, but not in exon 2 (Fig. 4c). At the same time, the maximal strength of mutation effects, measured as *d*_mut_/*d*, is similar for both exons 2 and 3, suggesting that, in principle, strong mutation effects are possible in exon 2 as well, but they point into the wrong direction.

Interestingly, the angle *θ* characterizing the directionality of a mutation effect in competition space is highly conserved across cell types. In Fig. 4d, we compare these angles between NALM6 and HEK293 cells and find that the results are consistent across systems, for both exon 2 (*r* = 0.71) and exon 3 (*r* = 0.95). On the other hand, the magnitude of the mutation vectors is much more variable between HEK293 and NALM6 cells, with low correlation coefficients, *r* = 0.10 and *r* = 0.16 for exons 2 and 3, respectively. Thus, whether a mutation affects the canonical splice site or its competing site, or both, appears to be hardwired in the architecture of the exon sequence (here the splice-site distance), and therefore tends to be conserved across cell types. In contrast, the strength of a mutation effect on spliceosome binding strongly depends on the cellular context, likely the expression of RNA-binding proteins, and is more variable across cell types.

To understand the different potential of exons 2 and 3 to be activated by single mutations, it is instructive to consider the sequence distance between the can-3’SS and alt-3’SS (Fig. 4e): In exon 3, the two splice sites are only 33 nucleotides (nt) apart from each other, whereas the distance is much larger in exon 2 (182 nt). Consequently, exon 2 mutations may not be able to simultaneously affect both splice sites, and therefore mutations point either vertically or horizontally in the two-dimensional competition landscape, independently affecting either the canonical or the alternative site (Fig. 3a). In contrast, in exon 3, the splice sites are so close to each other that a mutation frequently affects both splice sites, especially if it is located between the two splice sites. Then, a mutation typically weakens the upstream canonical site, while mildly strengthening the downstream site, implying that the mutation effect aligns with the direction of the optimal perturbation vector (Fig. 4a and c).

To test whether the same distance-dependence of mutation effects applies beyond alt-exon2 and alt-exon3, we analyzed other cryptic splice sites in the *CD19* minigene. We focused on 17 3’SSs in exon 3 and 13 5’SSs in exon 2 which harbor consensus splice-site motifs in the wildtype *CD19* sequence, AG for 3’SSs and GU for 5’SSs (Fig. 4f). Other cryptic splice sites lacking the consensus were ignored since they are generally weak and require multiple mutations for activation regardless of their positioning (Supplementary Fig. 7). For the chosen consensus splice sites, we related the distance of canonical and cryptic splice sites to the exon variant activation potential, measured as the maximal single mutation effect on exon variant inclusion in the NALM6 high-throughput mutagenesis results. In line with our hypothesis, we found that cryptic exon variants harboring a splice site nearby the canonical one tended to be stronger activated by single mutations when compared to exon variants spliced distally from the canonical version (Fig. 4f).

The distance-dependent alternative splicing patterns in *CD19* imply joint regulation of nearby splice sites and independent regulation of distant splice sites. To test whether such distance-dependent coordination of splice sites is a general phenomenon, we analyzed transcriptome-wide RNA-seq data for HeLa and K562 cells. Alternative 3’ and 5’ splicing events were quantified using MAJIQ^31,32^. The resulting dataset comprised thousands of alternative 3’ and 5’ local splicing variants (LSVs) in which a proximal (closer) and distal (further away) splice site were identified with respect to the reference donor exon (Supplementary Fig. 7b). The relative utilization of the proximal and distal splice site pairs was quantified using the percent spliced-in metric (PSI_prox_ = proximal/(proximal+distal)), ranging from 0 to 1 for each LSV. To exclude that alternative splicesite usage is dominated by splice-site strength, we considered only pairs of alternative splice sites with comparable splice-site strength expressed (MaxEnt score^33^, see Methods). This resulted in 1613 (837 3’ and 776 5’SS usage) and 2781 (1398 3’ and 1383 5’SS usage) events in HeLa and K562 cells, respectively. In line with the *CD19* results, splice site distance strongly influences the relative usage of proximal and distal 3’ splice sites in HeLa cells (Fig. 4g, blue line): when the competing distal splice site is very close (< 10 nt), the proximal 3’ splice site is preferentially used. Interestingly, this pattern is inverted at larger distances (10 – 30 nt), until the proximal splice site regains its prevalence at ∼ 40 nt with a strong PSI peak (∼ 75%). These fluctuations of splice-site choice support complex and concerted regulation mechanisms for nearby 3’ splice sites. Importantly, at longer distances (> 100 nt) the usage of proximal and distal 3’ splice sites is balanced, both fluctuating around PSI = 50%. Thus, in agreement with the *CD19* data (Fig. 4d), alternative 3’ splice sites further apart than 100 nt sites tend to be independently regulated in the genome-wide data. In further support for our findings in the *CD19* minigene, alternative 5’ splice site choice in HeLa cells showed a similar, but weaker, distance dependence compared to the 3’ splice sites, with minor shifts in turning point positions and amplitude (Fig. 4g, left, red line and points). Furthermore, in another cell line (K562), similar distance-dependent patterns of alternative 3’ and 5’ splice-site usage were observed (Fig. 4g, right).

Taken together, these data confirm that competing splice sites show uncoupled regulation if located more than ∼100 nt apart. At shorter distances, the relative usage of canonical and alternative splice sites depends in a complex way on their relative positioning.

### Pronounced directionality of mutation effects on alternative splice-site usage

Mutations in *CD19* exon 3 located in between the two competing splice sites had opposing effects on both variants, decreasing the recognition of the upstream canonical site while increasing that of the downstream alternative site (Figs. 4a and 3d). This suggests that *cis*-elements (e.g., changed by mutations) may exhibit a directional effect on splicing outcomes, differentially affecting upstream and downstream splice sites.

To further understand the coupled regulation of nearby splice sites, we turned to published mutagenesis datasets, in which the impact of cis-regulatory elements on multiple nearby splice sites was characterized across a large sequence space. To this end, we analyzed the published dataset by Rosenberg et al^16^ which determined the impact of two completely degenerate 25 nt regions on alternative splice-site usage, and therefore explored a large variety of sequences using a total of 265,137 minigene variants (Fig. 5a). Since the degenerate regions are located upstream and downstream of alternative 5’ splice sites in a single-intron minigene reporter, this dataset is ideally suited to test our hypothesis of directionality in splice-site regulation.

**Fig. 5.**
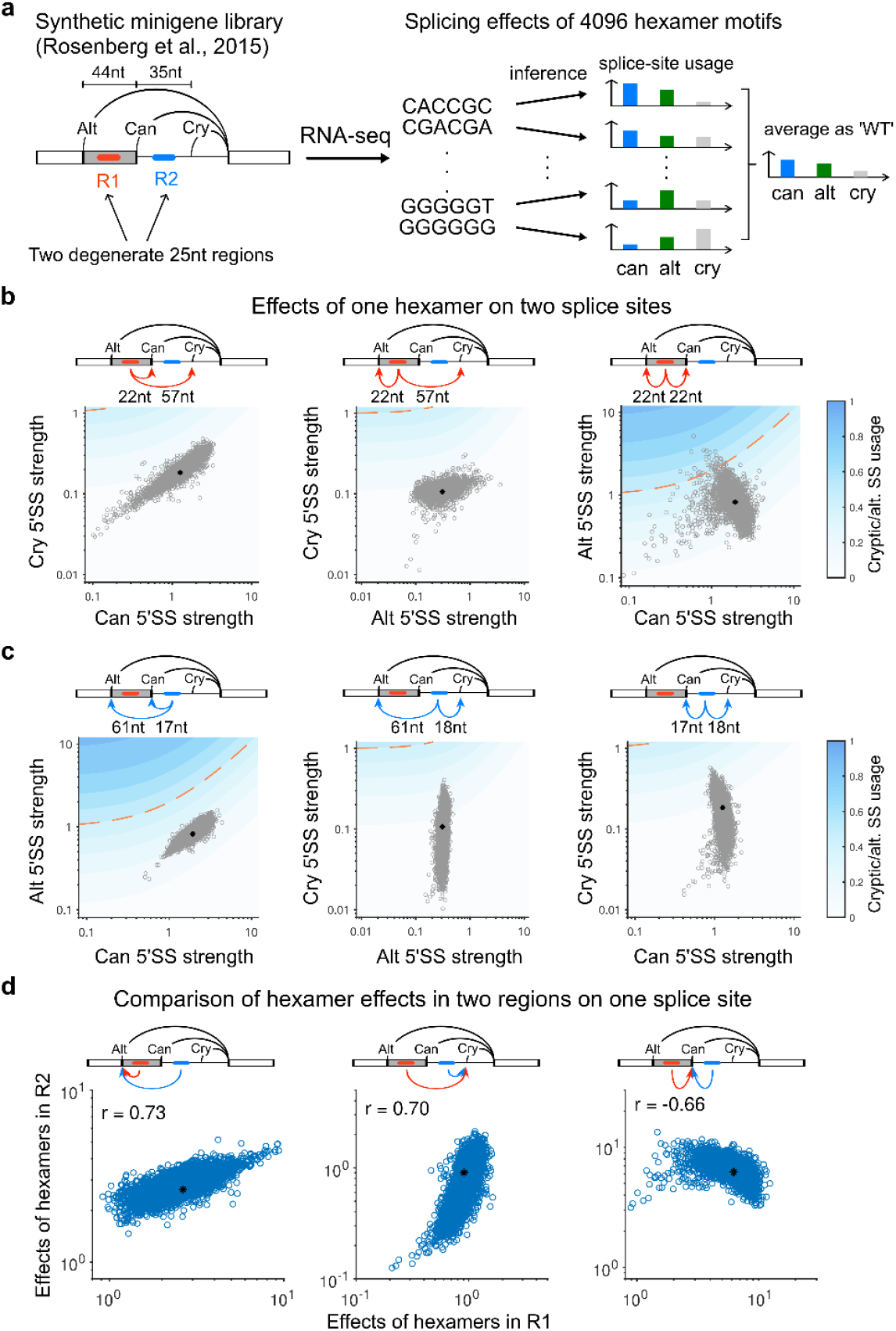
Mutations control upstream and downstream splice sites with opposite directionality. **a** Schematic representation of synthetic single-intron splicing system developed by Rosenberg et al.^16^, in which two degenerate 25nt regions (R1 and R2) were introduced to probe the sequence determinants of 5’ alternative splice site choice in a set of 265,137 minigene variants. RNAseq was used to quantify shifts from the canonical sites (can) to others, which are either typically present in the library at low frequencies (Alt: alternative splice site) or are absent (Cry: cryptic splice site). As *cis*-regulatory elements, 4096 hexamer motifs were considered. To quantify hexamer effects, the mean canonical, alternative and cryptic splice usage across all minigenes harboring that hexamer was compared to a hypothetical wildtype, measured as the mean splice site usage of all sequences in the library. **b – c** Coordinated effects of one hexamer located either to the exon (**b**) or the intron (**c**) on the usage of multiple nearby splice sites. Shown is the mean usage of the schematically indicated splice sites (x and y axis) for all minigenes containing a hexamer (grey dots) and for the wildtype (black dot), i.e., the mean of the complete minigene library. The splice sites are coordinately regulated when a hexamer is located upstream or downstream of both splice sites (first columns). When the motif is placed in between the two splice sites, it preferentially regulates the nearest one (middle columns). If the two splice sites are (approximately) symmetrically positioned around the motif, they tend to be antagonistically regulated (right columns). **d** Coordinated effect of the same hexamer motif located to distinct regions in the synthetic splicing reporter on the usage of a single target splice site. Shown is the mean usage of the schematically indicated splice site for all minigenes containing a given hexamer in R1 (x) and R2 (y), separately for each hexamer (blue dots) and for the wildtype (black dot). When both degenerate regions are positioned upstream or downstream of the target splice site the hexamer consistently enhances or inhibits splice site usage (left and middle panels). In contrast, the motif exerts opposite regulation when located upstream or downstream of the target splice site (right panel).

As the mutated 25 nt stretches are too degenerate to extract individual point mutation effects, we followed the approach by Rosenberg et al. and analyzed the effects of all 4096 possible hexamer motifs on alternative 5’ splice-site choice. For each hexamer, we calculated the average usage of the canonical, alternative and cryptic splice sites (Fig. 5a) in all minigenes containing that hexamer separately for each 25 nt degenerate region and compared this splicing outcome to the ‘wildtype’, defined as the average of all minigene variants in the screen.

Using this approach, we initially analyzed how a hexamer in a given degenerate region (25-mer) affects multiple nearby splice sites (Fig. 5b). As expected, we found that the hexamer coordinately regulates two splice sites that are either both located downstream of the motif (Fig. 5b, left) or both upstream of the motif (Fig. 5c, left). If the hexamer is located in between the two splice sites, the closer splice site seems to be primarily regulated, whereas more distant one is only weakly regulated (Fig. 5b, middle and Fig. 5c middle and right). If both splice sites are located with similar distance away from the degenerate region and show comparable strength, both tend to be simultaneously affected by the hexamer motifs (Fig. 5b, right). Interestingly, while coordinate regulation of splice sites occurs for a subset of motifs, the vast majority of hexamers regulates up- and downstream splice sites in opposite direction (Fig. 5c, right), similar to what we had observed for *CD19* exon 3. Hence, this supports that splice site regulation by sequence motifs indeed shows a strong directionality, with opposing effects on upstream and downstream splice sites.

To further support this, we also analyzed whether the same hexamer located to different 25-mers has distinct effects on a given target splice site. In line with our hypothesis, we observe that a hexamer located to a downstream 25-mer typically changes the usage of the target splice site in opposite direction when compared to the same hexamer in an upstream 25-mer (Fig. 5d, right). In contrast, the recognition of a splice site tends to be coordinately changed by a hexamer if both 25-mers are located downstream (Fig. 5d, left) or upstream (Fig. 5d, middle). Hence, a given hexamer located at different positions relative to a target splice site shows directional effects.

Taken together, we describe predictable effects of mutations nearby splice sites which may establish coordinated or anti-coordinated regulation of isoforms depending on their relative position. The results may lead to a better understanding of disease mutations leading to alternative splice-site choice.

## Discussion

The complexity of the human proteome is greatly enhanced by post-transcriptional and posttranslational modifications such as alternative polyadenylation, alternative splicing, RNA editing and covalent protein modifications. These modifications constitute a combinatorial code, leading to the generation of a wide variety of protein isoforms, especially in the human brain^1,3,34^. In this work, we show that kinetic models help to trace this high-dimensional combinatorial code back to a smaller number of molecular modification reactions, thereby providing insights into design principles and regulatory control.

Combining mathematical modeling and experimentation, we found that the complex alternative splicing outcomes of randomly mutated *CD19* minigenes are regulated by only two elementary rules: exon definition and competitive definition of exon variants with alternative use of 3’ and 5’ splice sites. Specifically, in *CD19* exons 1-3 the presence of respectively 7, 34 and 34 exon variants could theoretically lead to the generation of 9189 splice isoforms. By appropriate choice of 75 exon recognition parameters, our kinetic model could explain why only 93 isoforms are realized in the data, and at the same time it quantitatively accounted for all isoform frequencies and their shifts upon sequence mutations.

The model further provided insights into the context-dependence of mutation effects and simultaneously captured all major local splicing decisions, i.e., cassette exon skipping, alternative 3’ and 5’ splice site usage and intron retention. Previous work showed that mutation effects on splicing depend on the starting frequency of splice isoforms, and the term scaling law was coined for this phenomenon. According to the scaling law, the effect of a mutation (ΔPSI) on a binary splicing decision involving cassette exon skipping and inclusion depends on the starting isoform frequency (PSI) when characterized in different sequence backgrounds^14^. Our work is fully in line with these earlier reports but extends the scaling consideration from a one-dimensional projection to the full isoform space. Notably, even though we consider more dimensions, many of the mutation effects still follow the previously proposed scaling curve, showing maximal effects at intermediate isoform frequencies (Fig. 1b, Supplementary Fig. 1b-f). However, deviations occur if we analyze how a mutation affecting a particular splice site perturbs the genesis of splice isoforms employing competing splice sites (Fig. 1c). Such crosstalk between splice site mutations and splice isoform frequencies is difficult to comprehend intuitively. Therefore, mathematical models are valuable tools for the quantitative analysis of context-dependent mutation effects and highdimensional mutagenesis data. In fact, our models show that two simple rules, independent recognition of neighboring exons and mutually exclusive choice of competing alternative exon variants, are fully sufficient to represent the complete *CD19* dataset. Interestingly, the model is simple enough to directly infer the unknown splice site recognition parameters from isoform measurements by analytical calculations, implying that the framework does not require extensive computation and is therefore scalable to more complex splicing decisions.

To infer single mutation effects from measured combined mutation effects, we developed a hybrid approach, in which splice site recognition parameters inferred by kinetic modeling serve as an input for a linear regression model. Thereby, we could predict how single mutations affect splice site recognition and consequently splice isoform frequencies, providing mechanistic views of mutation effects on spliceosome assembly. In the future, such mechanistic insights may be derived for arbitrary sequence contexts by connecting the kinetic framework to sequence-based machine learning models trained on transcriptome-wide RNA-seq data. To achieve this goal, it will be necessary to represent splice junction reads arising from short-read sequencing by the kinetic model. Our current model can be readily accommodated to this scenario, since the currently modelled splice isoforms contribute to known junctions, i.e., a junction can be represented in the model by summing up the contributing isoforms. Therefore, without loss of generality, the modeling framework proposed here is widely applicable to publicly available genomic measurements to quantify, e.g., the genome-wide impact of RBP knockdowns on splicing decisions. Since the multi-dimensional footprint of a perturbation is much more informative than one-dimensional splicing metrics, we expect that the quantitative modeling of genome-wide RBP knockdown data as proposed here will lead to more detailed mechanistic insights into RBP action.

Following up on the inferred mutation effects on *CD19* isoforms, we could show that splice site distance is an important parameter determining whether the switch from canonical to alternative/cryptic splice site can be achieved by single mutations. Our data shows that a mutation in between two nearby competing splice sites simultaneously affects both in opposite direction, implying a pronounced directionality of mutation effects on up- and downstream splice sites which decays with distance. Early reports, in which splicing enhancers and suppressors were placed at different distances to a splice site indeed showed a monotonic decay of splicing effects, with strongly weakened effects at distances exceeding 100 nt^31,34–39^ as observed for *CD19* (Fig. 4f). Our analysis of genome-wide splicing datasets confirms uncoupling of distant splice sites, as we do not observe preferential use of proximal vs. distal splice sites at distances beyond ∼100 nt in both analyzed cell lines (Fig. 4g). For nearby splice sites, we find that the first (proximal) 3’ splice site in the intron is commonly favored at low distances from 3 to 10 nt. This observation is in line with earlier findings showing that the spliceosome almost exclusively chooses the proximal splice site (i.e., the closest one to the branch point and poly-pyrimidine tract) for short SS distances^36,37^. Published work further showed an enrichment of cryptic 3’ splice sites about 12 to 21 nucleotides upstream of canonical splice sites^38^. Since these proximal cryptic splice sites are by definition only weakly utilized, this may explain why we observe preferential distal splice site usage in this distance window (Fig. 4g). In conclusion, distance-dependent effects on alternative splice choice may reflect splice site positioning relative to core sequence elements such as the branchpoint and the poly-pyrimidine tract. The absence of these core sequence elements in the alternative 5’ splice site architecture may explain why these generally show weaker distance-dependent splice site coupling effects (Fig. 4g).

Taken together, our work shows that kinetic models help to trace back coordinated splicing changes in high-dimensional RNA sequencing data to a limited number of molecular events at the splice site level. When applied to data from patients, kinetic modeling approaches may lead to mechanistic insights into disease mechanisms and may aid the design of splicing therapies, in which antisense oligonucleotides target specific splice sites or nearby *cis*-regulatory elements.

## Acknowledgements

We thank Stefanie Ebersberger and Mario Keller for the initial processing and analysis of the transcriptome-wide RNA-seq data on splice-site choice. We thank Sylvia Weiss and Daniela Ramesohl for their technical assistance with experimental work. This work was funded by the Deutsche Forschungsgemeinschaft (DFG) to K.Z., J.K. and S. L. (ZA 881/2–3 to K. Z., KO 4566/4-3 to J. K., and LE 3473/2–3 to S. L.). C.L. and S.L acknowledge support by the Stuttgart Research Center Systems Biology (SRCSB).

## Author contributions

Conceptualization, C.L., S.U. and S.L.; Data Curation, C.L. and S.U.; Formal analysis, C.L., S.U., M.K., M.B. and S.L.; Investigation, C.L., S.U., M.K. and M.B.; Methodology, C.L., S.U., M.K., K.Z., J.K. and S.L.; Validation: C.L. and S.U.; Resources, K.Z., J.K and S.L.; Software, C.L., S.U., M.B. and S.L.; Visualization, C.L., S.U. and S.L.; Supervision, C.L. and S.L.; Writing – original draft, C.L., S.U. and S.L.; Writing – review & editing, C.L., S.U., K.Z., J.K., and S.L.

## Methods

### A kinetic model of splice-site competition

To quantitatively understand how splice-site competition gives rise to different RNA isoforms, we studied a minimal model where two possible 3’SSs, canonical (can-3’SS) and alternative (alt-3’SS), can be used in an exon that is flanked by two constitutive exons (Fig. 1a). We assumed that the selection of the can-3’SS and alt-3’SS is determined by exon definition mechanism, in which the two sites compete for the same downstream 5’SS in forming a cross-exon complex with the early spliceosome (U1 and U2). The competitive exon definition gives rise to three types of isoforms – inclusion of the canonical and alternative exon variants (can-exon and alt-exon), respectively, as well as the skipping isoform in which the whole exon is skipped (Fig. 1a). We assumed that the spliceosome binding is reversible and much faster than the RNA synthesis and splicing catalysis, and thus the spliceosome binding states determine the splicing outcomes. The dynamic transitions of the binding states can be described by the following ordinary differential equations (ODEs).

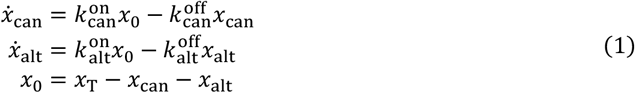

where *x*_can_ and *x*_alt_ are the levels of the transcripts with U1-U2 bound cross the can-exon and alt-exon variants, respectively. *x*_0_ represents the abundance of the unbound state and *x*_T_ the sum of all three binding states. 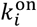 and 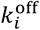 (*i* = can, alt) are the association and dissociation rate constants of U1-U2 assembly cross an exon variant (can for canonical and alt for alternative exon variant). Based on the fast-binding assumption, we derived the solution of Equation 1 at steady state.

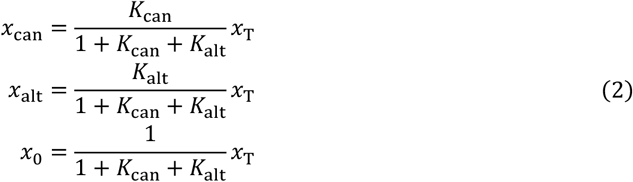

where 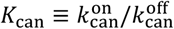 and 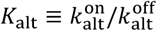 are the equilibrium constants of spliceosome binding to the can-exon and alt-exon, respectively. The distribution of the three binding states is thus determined by the affinities of early spliceosome assembly across different exon variants. We further assumed that the three spliceosome-binding states are processed with the same splicing catalysis rate and that the three resulting splice isoforms share the same degradation rate. Then the fractions of the binding states predominantly determine the frequencies of the splicing products,

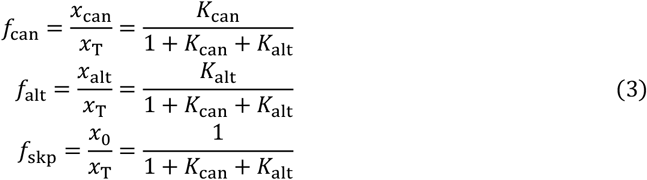

where *f*_can_ and *f*_alt_ are the frequencies of isoforms including the can-exon and alt-exon variants, respectively, and *f*_skp_ is the frequency of the exon skipping isoform. *f*_can_ and *f*_alt_ can also be interpreted as recognition probabilities of the can-exon and alt-exon variants, respectively, and *f*_skp_ = 1 - *f*_can_ - *f*_alt_ is then the probability of the whole exon not being recognized (or defined). Thus, the isoform frequencies visualized in Fig. 1a are jointly shaped by the affinities of early spliceosome binding to two exon variants.

### Maximum-likelihood inference of overall splicing isoforms of *CD19* minigene

To quantitatively explain the extensive usage of alternative/cryptic 3’ and 5’ splice sites in *CD19* minigene splicing, we devised a two-tiered modeling approach. In a first coarse-grained modeling step, we extended the above-described modeling approach and described splicing decisions towards three major splicing fates, i.e., exon 2 inclusion, exon 2 skipping and second intron retention. In a second fine-grained modeling step, we refine this description and additionally consider the local splice site choice in each exon. Thereby, we derive a quantitative and predictive model describing the abundance of overall 93 splice isoforms.

For coarse-grained modeling, we ignored the details in alternative splice site usage in each exon and considered lumped pools of exon 2 inclusion, exon 2 skipping and second intron retention. We employed a previously developed kinetic model^28^ which assumes a so-called exon definition mechanism, in which the spliceosome initially recognizes (defines) the exons as functional units. Then, splicing decisions are made based on the combination of exons that are recognized in a transcript, assuming that an intron can only be spliced if both neighboring exons are defined (Fig. 1a). In quantitative terms, the initial exon definition step is described by the spliceosome binding to each exon, and the occupancy can be interpreted as exon recognition probability, denoted as *α, β* and *γ* for exon 1 – 3, respectively. The exon recognition probability is a function of the spliceosome binding affinity as the following,

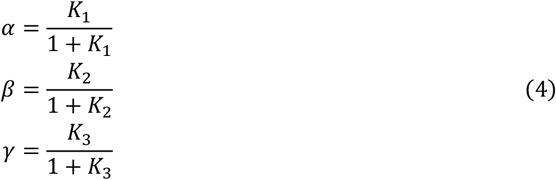

where *K*_1_, *K*_2_ and *K*_3_ are the equilibrium constants of spliceosome binding to exon 1, 2 and 3, respectively. The spliceosome binding affinity controls how likely an exon is defined. For instance, if the spliceosome binding is extremely weak (*K*_1_→0), then exon 1 will not be defined (*α* = 0); strong binding (very large *K*_1_) results in full recognition of the exon (*α* = 1). Importantly, in our coarse-grained model, we do not distinguish which exon variant is chosen but only describe whether any of the exon is defined or not.

To link spliceosome binding to splicing fates, we assume that the spliceosome assembles independently at each exon. Thus, the binding states of the minigenes are combinations of defined (denoted by 1) or/and undefined (denoted by 0) exons 1 – 3. Following exon definition mechanism, states 111 (all exons recognized), 101 (middle exon not recognized) and 110 (last exon not recognized) give rise to inclusion, skipping and second intron retention isoforms, respectively. The corresponding joint probabilities are calculated based on the exon recognition probabilities and are given by,

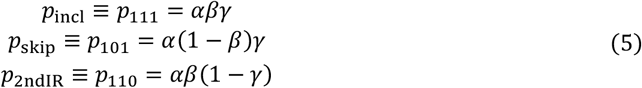

To infer the recognition parameters (*α, β* and *γ*) from the *CD19* mutagenesis data, we constructed a likelihood function linking the modeled isoform probabilities in Equation 5 and the counts of the three isoforms *n*_incl_, *n*_skip_ and *n*_2ndIR_ (inclusion, skipping and intron 2 retention, respectively) recorded by RNA-seq,

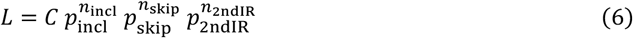

where *C* is a multinomial coefficient that measures the number of all possible microstates of isoform counts:

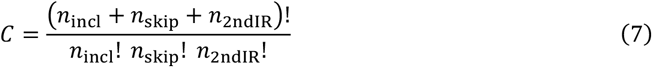

The likelihood function lacks the probability terms for the first and full intron retention isoforms, because neither was found in the splicing products of WT and mutated *CD19* minigenes. Combining Equations 5 and 6, the likelihood of observing the data is a function of the three model parameters (*α, β* and *γ*):

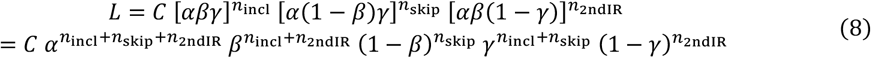

Our goal is to find the parameter values which maximize the likelihood function of the current observation (*n*_incl_, *n*_skip_ and *n*_2ndIR_). For convenience, we transform the likelihood into logarithmic scale,

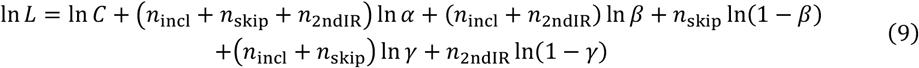

Maximization of log-likelihood is equivalent to maximizing the likelihood function. The first term in the equation is a constant that is independent of the model parameters. The three parameters contribute to the log-likelihood function independently. The parameter *α* is different from the other two parameters, as it monotonically controls the likelihood value. *α* maximizes the likelihood when it reaches its largest possible value. By definition *α* is the recognition probability or U1-U2 complex occupancy of the first exon, so its maximum value is 1,

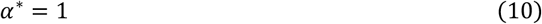

where the superscript “*” represents the optimal parameter value that maximizes the likelihood. This result suggests that the first *CD19* exon is always successfully defined, which is consistent with the fact that no first or full intron retention isoform was detected.

For the other two parameters *β* and *γ*, the log-likelihood function will only peak at an intermediate value between 0 and 1. The necessary condition for reaching the peak is that the slopes of the log-likelihood are zero,

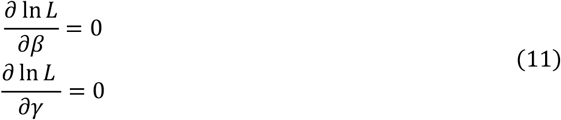

The equations can be solved analytically to yield the optimal values of *β* and *γ*,

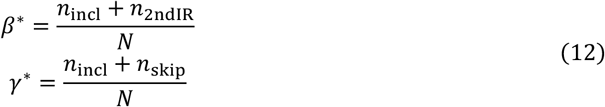

where *N* is the sum of all the isoform counts. The analytical result suggests that the best fit exon recognition frequency is the fraction of all isoforms that contain the corresponding successfully recognized exon. Substituting the above results (Equation 12) into Equation 4, the affinities of spliceosome binding to exon 2 and 3 (*K*_2_ and *K*_3_) can be calculated.

Using the best-fit parameter values, we successfully reproduced the inclusion, skipping and second intron retention frequencies observed in ∼ 9000 minigenes (Supplementary Fig. 3) and confirmed that the other possible isoforms (first intron and full intron retention) that were not detected experimentally are indeed in very low abundance (Supplementary Fig. 3). Thus, the simple exon-definition model successfully captures the overall splicing outcomes of *CD19* minigenes. The successful model fitting suggests that the three exons in the minigene are defined independently of one another and that *CD19* splicing decisions depend on the definition of exons as relevant functional units.

### Model extension and maximum-likelihood inference of alternative splice-site usage in *CD19* minigenes

The large-scale mutagenesis experiments on *CD19* minigene revealed extensive activation of alternative and cryptic splice sites, especially in exon 2 and 3 (Fig. 2a). To test whether the maximum likelihood method can also be applied to infer the generation of a large number of complex isoforms using alternative/cryptic splice sites, we extend the coarse-grained exon-definition model considering the details of alternative splice-site choice in each exon. In addition to pairing the canonical 3’ and 5’ splice sites for exon definition, we here also allow the recognition of many exon variants defined by different alternative/cryptic 3’ and 5’ splice-site pairs. For *CD19* minigene, exon 1 has 7 possible 5’SSs (including the canonical site) and thus 7 exon variants. In exon 2, 6 3’SSs and 24 5’SSs combine into 34 detected exon 2 variants. 34 5’SSs in exon 3 give rise to 34 exon variants. In line with the previous simple model, the initial binding of spliceosome dominates the splicing decision. With multiple variants of the same exon, they are assumed to compete for spliceosome binding, and therefore the recognition of these exon variants is mutually exclusive. The overall recognition probability of a whole exon, which is quantified in the previous section (Equation 4), is then the sum of the recognition probabilities of all variants of that exon:

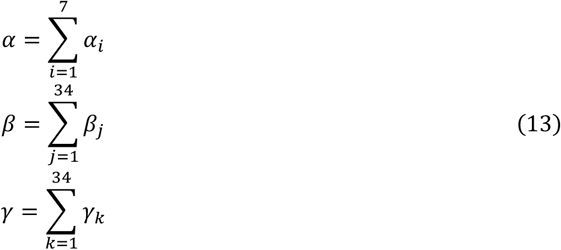

where *α*_*i*_, *β*_*j*_ and *γ*_*k*_ are the recognition probabilities for exon variants of exon 1, 2 and 3, respectively. The recognition probability of each exon variant is determined by how strongly the spliceosome binds to that variant in competition with other variants of the same exon, and it is thus derived assuming competitive spliceosome binding,

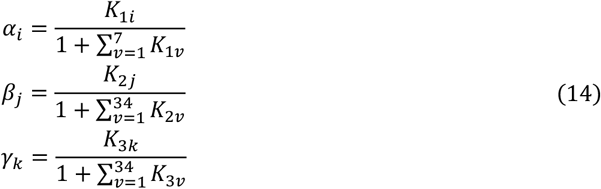

where *K*_*xv*_ represents the affinity of the spliceosome binding (*k*_on_/*k*_off_) to the *v-*th variant of exon *x* (*x* = 1, 2, 3).

In general, assuming independent combination of these exon variants, the model describes 8092 (7 × 34 × 34) inclusion and 238 (7 × 34) skipping isoform variants of *CD19* minigene. For the first intron retention isoform variants, all possible 5’SSs in exon 1 and 3’SSs in exon 2 are degenerate, so the pairing between 5’SSs in exon 2 (24) and 3’SSs in exon 3 (34) yields 816 (24 × 34) first intron retention isoform variants. Likewise, there are 42 (7 5’SSs in exon 1 × 6 3’SSs in exon2) possible second intron retention variants. Including one full intron retention isoform (intact premRNA without splicing), all exon variants of *CD19* can constitute in total 9189 possible isoforms.

Using RNA-seq, we only detected about 1% (93 out of 9189) of the estimated *CD19* isoform variants that fall into only three classes: inclusion, skipping and second intron retention. To describe these three isoform categories, we assume the independent combination of exon variants. In similarity to Equation 5, any inclusion isoform variant is produced by independently selecting an exon variant in each exon, and its probability can be quantified as,

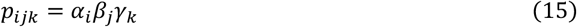

Likewise, the frequency of a skipping isoform variant, where none of the exon 2 variants is selected, is then,

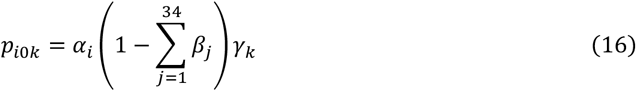

In second intron retention, only the 3’ splice site but not the 5’ splice site of exon 2 is spliced. In the initial exon recognition step, it is therefore only relevant that the correct 3’ splice site is recognized, whereas the 5’ splice site is degenerate. Hence, the exon 2 recognition probability that contributes to second intron retention is the sum of all exon-variant probabilities using a common 3’ splice site under consideration. This is indicated as:

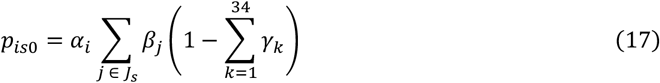

where *J*_*s*_ is an ensemble of indices representing all alternative exons that share the same 3’ splice site *s* in exon 2.

To fit the extended model to the full dataset of 93 isoforms found in ∼ 9000 *CD19* minigene variants, we constructed the likelihood function in the same vein as in Equation 6,

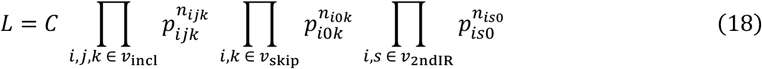

where *v*_incl_, *v*_skip_ and *v*_2ndIR_ are sets of indices (*i, j, k* and *s*) marking the inclusion, skipping and second intron retention isoform variants, respectively. The constant *C* is the combinatorial count of the current observation, similar to that in the simple model:

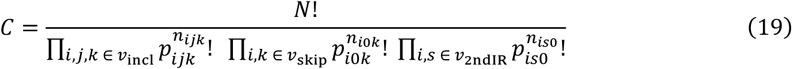

where *N* is the total count of all isoforms. To facilitate mathematical derivation, we transformed the likelihood function into to logarithmic scale,

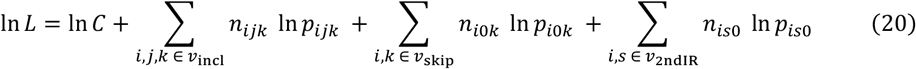

The first constant term does not contribute to finding the parameter values that maximize the likelihood. Using the same technique as in the previous section, the necessary condition of loglikelihood reaching maximum is:

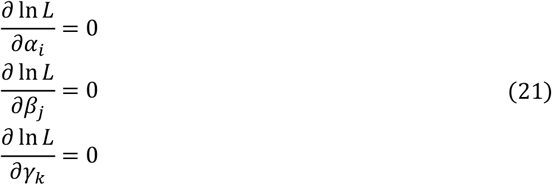

where *i* = 1, …, 7 and *j, k* = 1, …, 34. Interestingly, for such a complex system of alternative splice site choice, analytical solution still exists. The exon recognition probabilities of the outer exon (exon 1 and 3) variants are the fractions of isoforms that contain the relevant exon variants:

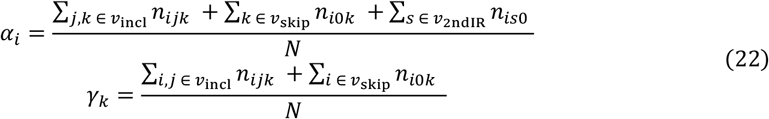

The intron 2 retention isoforms contain alternative middle exons that use a common 3’ splice site and degenerate 5’ splice sites, so the recognition probability of the alternative middle exon *j* reads:

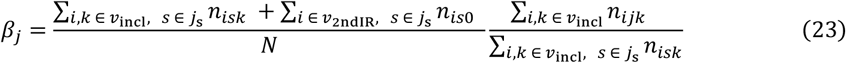

This is a product of two fraction terms: the first term is the fraction of all isoforms (inclusion and retention isoforms) that utilize the common 3’ splice site of the alternative middle exon *j*; the second term is the fraction of the exon variant *j* in all inclusion isoforms that use the common 3’ splice site *s* (*j*_s_ in the summation index).

Using the analytical solutions of *α*_*i*_, *β*_*j*_ and *γ*_*k*_ (*i* = 1, …, 7 and *j, k* = 1, …, 34), we reproduced frequencies of all 93 isoforms in ∼ 9000 *CD19* minigenes (Supplementary Fig. 2). Therefore, the generation of complex alternative isoforms can be quantitatively described by two simple principles – exon definition mechanism and mutual exclusive definition of exon variants.

### Inference of single-mutation effects on *CD19* minigene splicing by regression analysis

To learn the single-mutation effects on recognition of all 75 exon variants and the 93 resulting isoforms, we assumed that the mutation effects are additive at level of spliceosome binding free energy. The equilibrium constant of an exon variant for a given minigene *i* with specific mutations can be described by,

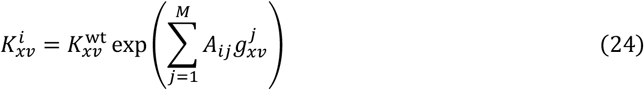

where 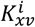 is the spliceosome affinity to the *v-*th variant of exon *x* in minigene *i*, and 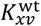 is the wildtype counterpart without mutations. 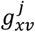 represents the effect of a single mutation *j* on the binding free energy and A_*ij*_ is a binary (0 or 1) value showing whether the mutation *j* occurs in minigene *i*. We found in total 4484 unique single mutations (*M* = 4484; 3591 single nucleotide substitutions and 893 nucleotide insertion or deletion) in 8996 minigenes from replicate 1 and 4460 mutations (3571 single nucleotide substitutions and 889 nucleotide insertion or deletion) in 8597 minigenes from replicate 2.

Taking logarithms on both sides of Equation 24, we obtained a linear equation system.

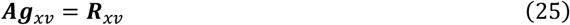

where ***A*** is an M (number of single mutations) by *N* (number of minigenes) binary matrix indicating the occurrence of mutations in the minigenes (0 representing absence and 1 presence). ***g***_*xv*_ is a 3591 (number of single mutations) by 75 (number of exon variants) matrix quantifying the single mutation effects for a given exon variant (*v-*th variant of exon *x*), which is to be inferred by model fitting to data. And 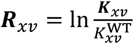 is the log ratio between equilibrium constants of the minigenes and that of the wildtype for the *v-*th variant of exon *x*, which can be directly calculated from data using the maximum likelihood method in the previous section. Therefore, the inference of single mutation effects can be reduced to solving linear equation systems by linear regression methods for each of the 75 exon variants. To avoid overfitting due to the large number of mutations, we applied L1 regularization (Lasso) in the regression to remove unimportant mutations. 10-fold cross validation was used to optimize the penalization strength and test predictive power of the penalized regression (Fig. 2d).

To predict the single mutation effects on the abundance of 93 isoforms (Supplementary Fig. 4), the spliceosome affinities to the canonical and alternative/cryptic exon variants upon single mutation *m* can be calculated (in a similar way for the minigenes by Equation 24),

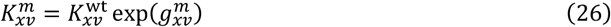

And the corresponding usage of each exon variant upon a single mutation *m* can be calculated by substituting Equation 26 into Equation 14,

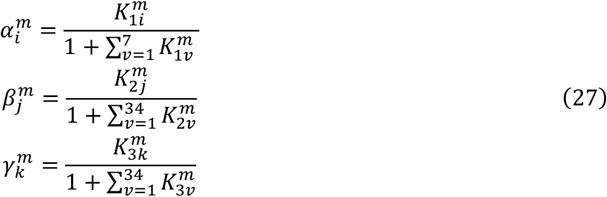

Using Equations 15 – 17, the frequencies of the 93 isoforms of a hypothetic *CD19* single mutant (with mutation *m*) can be calculated (Supplementary Fig. 4),

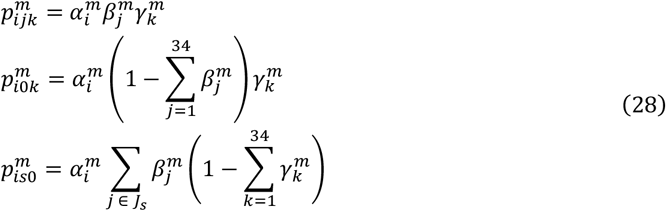

where 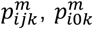 and 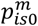 are the frequencies of inclusion, skipping and second intron retention isoform variants for the *CD19* minigene with a single mutation *m*. Consistent with Equations 15 – 17, the indices (*i, j, k* and *s*) indicate the combination of different variants in exon 1 – 3.

To further study how a specific alternative/cryptic exon variant is activated by competing with the canonical exon, we mapped the single mutation effects on both exon variants to the 2D splicesite competition landscape (Fig. 1). We focused on the activation of alt-exon2 variant (using alt-3’SS at position 658) of exon 2 and alt-exon3 variant (using alt-3’SS at position 1073) of exon 3. To map multiple different single mutations onto the same 2D landscape (as in Fig. 3a and Fig. 4a), we renormalized the inferred spliceosome affinities binding to the canonical and alt-exon2 (or alt-exon3) variant by the sum of affinities of all other cryptic exon variants. For instance, we describe the competition between the canonical exon 2 and alt-exon2 by expanding Equation 27 (for *β*),

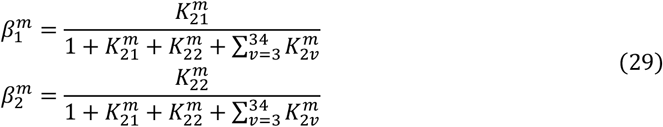

where 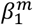 and 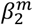 are the recognition probabilities of the canonical exon 2 (*j* = 1) and alt-exon2 (*j* = 2) upon mutation *m*, respectively. 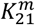 and 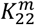 are the corresponding spliceosome binding affinities to the canonical and alt-exon2 variants, respectively, while 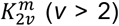 represent the affinities for the rest cryptic exon 2 variants. Using a common normalization factor 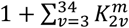, Equation 29 reads,

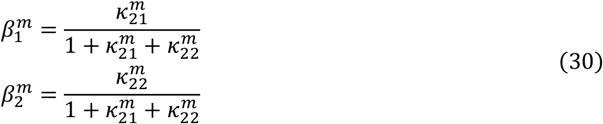

where

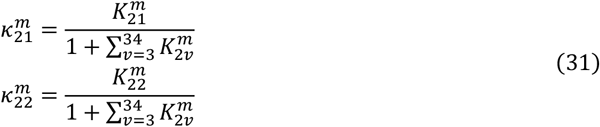

The parameters 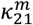 and 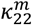 are the apparent definition strengths of the canonical exon 2 and alt-exon2 upon mutation *m*, respectively. For WT *CD19* minigene, it follows the same mathematical form replacing *m* by WT. The derived Equation 30 has the same form as Equation 3 in which only two exon variants are possible. Therefore, we use Equation 30 to map the high dimensional exon variant space to a 2D competition landscape. Note that the renormalization does not change the key features, i.e., the movement in the landscape, of the mutation effects, because all other cryptic exon variants show very weak recognition probabilities (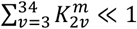in Equation 31). It is possible that a mutation strongly activates another cryptic splice site in exon 2 (increasing 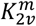) which reduces both 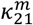 and 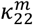 (according to Equation 31). In this case, the mutation does not directly weaken the recognition of canonical exon 2 and alt-exon2, so the resulting splice product interpreted from the 2D landscape viewpoint should be adjusted. The product is not the exon 2 skipping isoform, as would be predicted from Equation 3, but a cryptic inclusion isoform variant using neither the canonical exon nor alt-exon2. However, this is a rare event, as most mutations simultaneously reducing 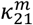 and 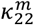 give rise to the exon 2 skipping isoform (Supplementary Fig. 4).

In the same vein, we can also map the high dimensional exon recognition parameter space to 2D landscape for competition between canonical exon 3 and alt-exon3, using the following equations,

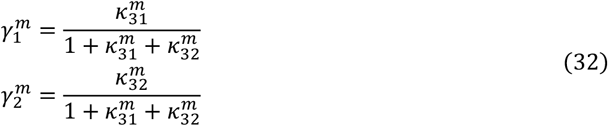

where 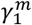 and 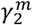 are the recognition probabilities of the canonical exon 3 and alt-exon2 upon mutation *m*, respectively. The parameters 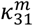 and 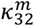 are the corresponding apparent definition strengths, defined by,

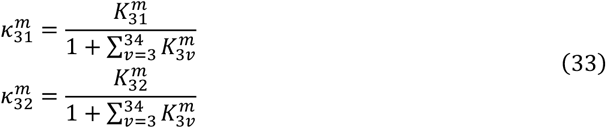

Using Equations 30 – 33, we quantified and characterized the effects of single mutations in the context of splice-site (or exon variant) competition in Fig. 3 and Fig. 4.

### Validation of additive mutation effects

To test the single mutation effects inferred from the linear regression method, we constructed 40 *CD19* minigene variants harboring single mutations and transfected them into HEK293 cells. Using capillary electrophoresis, we measured the frequencies of the five major isoforms: exon 2 inclusion, skipping, intron 2 retention, alt-exon2 and alt-exon3. These results well support our model predictions (Fig. 2f and Supplementary Fig. 5e). Next, we sought to test our linear regression assumption that the mutation effects are independent and additive (Equation 24). To this end, we produced 15 double mutants pairing some of the above tested single mutations located in exon 2 and measured their splicing outcomes using the same methods for the single mutants.

To quantitatively model these double mutants, we reduced the high-dimensional exon definition model (Equations 13 – 17) by considering the generation of four isoforms – inclusion, skipping, intron 2 retention and alt-exon2, as mutations residing in exon 2 do not activate alt-exon3. We further assume that the mutation effects add up at the spliceosome binding energy level (in line with Equation 24). Specifically, the exon recognition probabilities of the WT *CD19* read,

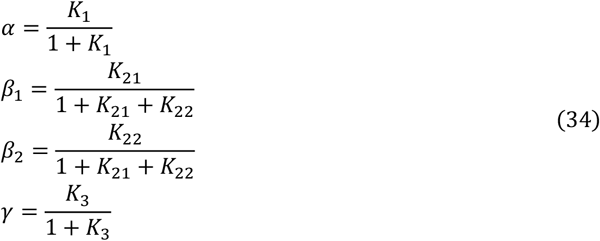

where *α, β*_1_, *β*_2_ and *γ* are the recognition probabilities of exon 1, canonical exon 2, alt-exon2 and exon 3, respectively; and *K*_1_, *K*_21_, *K*_22_ and *K*_3_ are the affinities (equilibrium constant) of spliceosome binding to the corresponding exons or exon variants. The splice isoforms can then be quantified by,

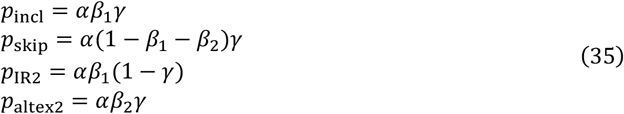

where *p*_incl_, *p*_skip_, *p*_IR2_ and *p*_altex2_ are the frequencies of inclusion, skipping, intron 2 retention and alt-exon2 isoforms.

We assume that the mutations in exon 2 only affect the recognition of exon 2 variants (canonical and alt-exon2) but not exon 1 and 3. A single mutation A in exon 2 can modulate the spliceosome binding to both exon variants,

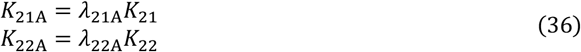

where *λ*_21A_ and *λ*_22A_ are the fold changes of spliceosome binding affinities to canonical and alt-exon2 variants upon mutation A, respectively. Similarly, for single mutation B, we have,

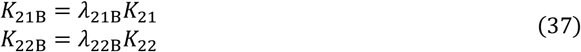

where *λ*_21B_ and *λ*_22B_ are fold changes in recognition strengths for the two exon 2 variants. If the double mutations A and B occur, the spliceosome binding energy change is the sum of those caused by individual mutations. Following Boltzmann distribution, i.e., fold change ∝ Exp(free energy change), the logarithmic fold changes are additive, which is described as,

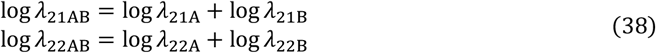

And this logarithmic additivity leads to multiplicative relation for the fold change parameters,

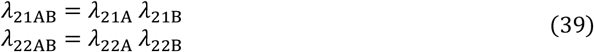

Using Equations 36 – 39, the probabilities of exon 2 variants upon single (*β*_*i*A_ and *β*_*i*B_) and double (*β*_*i*AB_) mutations can be derived (*i* = 1, 2; 1: canonical exon2; 2: alt-exon2),

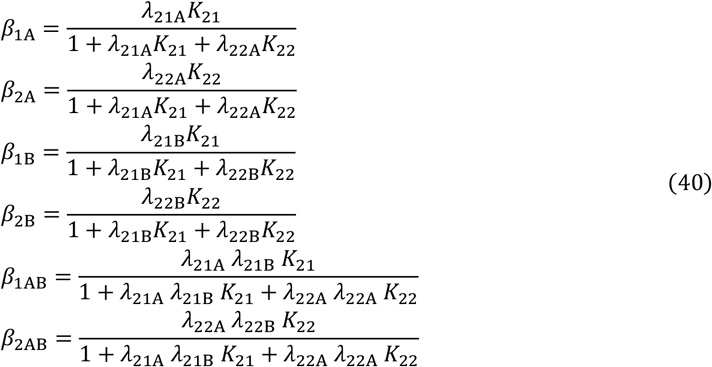

The mutant isoform frequencies are described in similarity to Equation 35,

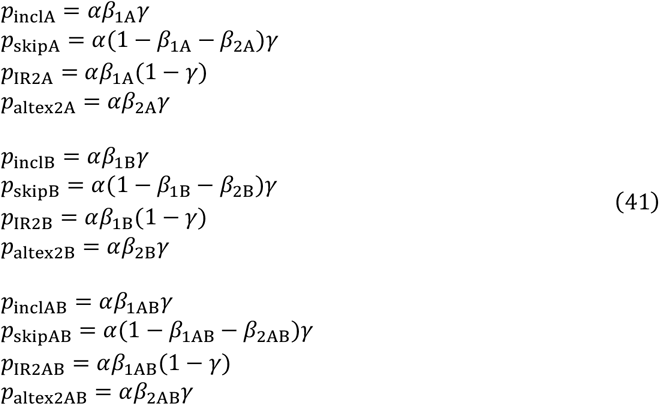

where *p*_*X*A_, *p*_*X*B_ and *p*_*X*AB_ are the frequencies of the splice isoforms (*X* = incl, skip, IR2 and altex2) for single mutants A, B and double mutants AB, respectively.

To challenge the assumption of additive mutation effects, we fit the model (Equations 35 and 41) jointly to the WT, single and double mutation data using maximal likelihood method (similar to Equation 6). The likelihood function for the combined mutations reads,

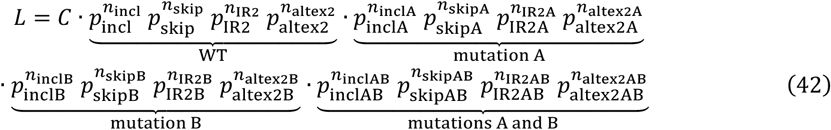

where *n*_*X*_, *n*_*X*A_, *n*_*X*B_ and *n*_*X*AB_ are the measured abundance of splice isoform *X* (incl, skip, IR2 and altex2) for WT, single mutants A, B and double mutant AB *CD19* minigenes, respectively. *C* is a constant combinatorial number.

Using numerical optimization approach (gradient descend method), we minimized the −2 log *L*, which is equivalent to maximizing *L*, to obtain the fold change parameter values (*λ*s). We then simulated the isoform frequencies for the single and double mutants using the best-fit parameters and found the simulation results agree very well with the data (Fig. 3b and Supplementary Fig. 6), indicating the validity of the mutation-effect additivity.

### Analysis of MAJIQ/modulizer data for alternative 3’ and 5’ events

To investigate alternative splice site choice on a genome-wide scale, we analyzed two datasets in which global splicing patterns were quantified from RNAseq-data using the splicing quantification tool MAJIQ^31,32^. MAJIQ quantifies transcriptome-wide splicing in the form of local splicing variants (LSVs) and these LSVs cover all detected splicing junctions connected to one reference exon (Supplementary Fig. 7b). The splicing outcome of each LSV is quantified using a PSI value, defined as the usage of each junction in the LSV relative to all other detected splicing events in that LSV (see MAJIQ documentary https://majiq.biociphers.org/majiq-spel/docs). For alternative 3’ or 5’ splice sites, MAJIQ operates with “proximal” and “distal” splice site pairs, based solely on the distance of the splice site to the reference exon (Supplementary Fig. 7b). In some LSVs, selection of a proximal/distal splice site was not the only set of options, but additional splicing events like intron-retention were detected as well. To concentrate our analysis on alternative splice site choice only, we normalized the PSI values of each pair of corresponding proximal/distal splice sites to 1 by:

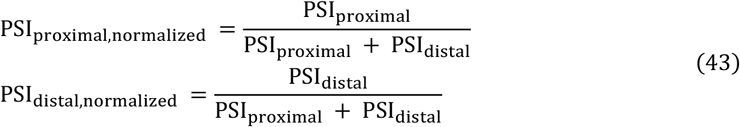

Consequently, this normalization yielded PSI_proximal,normalized_ + PSI_distal,normalized_ = 1.

In doing so, each pair of splice sites was treated as a binary event. Splice site distance, the distance between proximal and distal splice site (Supplementary Fig. 7b), was extracted from the modulizer files (termed “event size”, expressed in nt) for each splice site pair. For each dataset, and for alternative 3’ and 5’ events separately, PSI values of splice site pairs were sorted by splice site distance. Both distance and PSI values were then binned with bin size = 50 (unfiltered data, Supplementary Fig. 7c) or 20 (filtered data, Fig. 4g, see section MaxEnt^33^ filtering and data processing), and the median PSI and mean distance were calculated per data bin. The data were plotted using only the median PSI_proximal_ values of both 3’ and 5’ events since, due to normalization, PSI_proximal_ and PSI_distal_ were completely symmetrical around the value of 0.5. A running mean (window size 3 to 5) was added to highlight tendencies of data distribution.

### MaxEnt filtering and data processing

To exclude PSI pairs in which the preferential use of proximal or distal splice site was mainly caused by different splice site strengths masking distance/intron length effects, the modulizer data were filtered for splice site pairs, in which both splice sites had similar strength. Splice site strength was determined based on sequence using the Maximum Entropy model^33^. To do so the genomic sequences of each splice site were extracted from the human reference genome (hg38) using R and the public R libraries BSgenome and Biostrings. The splice site sequences were used to assemble the 9- and 23-mers required for the MaxEnt algorithm to calculate scores for 5’ and 3’ splice sites, respectively. Similarity of MaxEnt scores per splice site pair was assessed by calculating the difference between proximal and distal splice site scores and was defined as a ΔMaxEnt score within -2.5 and +2.5. All splice site pairs without similar MaxEnt scores were excluded, which accounted for about 60% of data points.

### Quantifying effects of hexamer motifs on splice-site choice from synthetic minigene libraries reported by Rosenberg et al

Rosenberg et al. screened an alternative 5’ splice site library of 265,137 minigenes consisting of one intron flanked by two exons^16^. An alternative splice site (SD1) is located upstream of the canonical site (SD2), both being in the exonic region. In addition, a cryptic splice site resides in the downstream intronic region. These splice sites are regulated by two degenerate 25-nt regions containing random sequences, locating upstream (R1) and downstream (R2) of the canonical splice site, respectively (Fig. 5a).

To infer how sequence contexts control splice-site choice, we followed the same approach as Rosenberg et al. and focused on the effect of 6-mer motifs. To quantify the effects of a 6-mer motif on the splice-site choice, for each of the 4096 possible 6-mer motifs, the isoform distribution was calculated by taking the mean isoform frequencies of all minigenes containing the motif. This calculation was done separately for the left and right degenerated region. To ensure stability of the results, only minigenes containing more than a total of 100 reads were considered.

Per definition, the dataset contains no wildtype construct. To nevertheless estimate the effect sizes of the motifs compared to a reference situation, we defined the ‘WT’ splice-site usage as the mean effect sizes of hexamers across both degenerative regions independent of the presence of motifs. To quantify the effect of hexamer motifs on splice-site choice, we applied our multidimensional model to describe the competition among the recognition of the canonical, alternative and cryptic 5’ splice sites. For the above defined ‘WT’, we modeled the frequency of splice-site usage as the following,

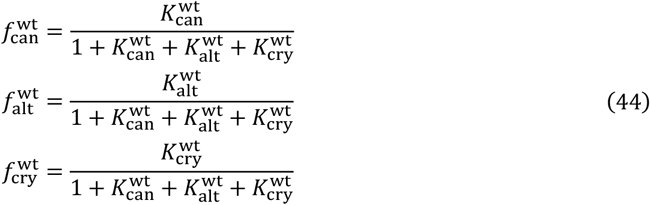

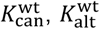 and 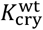 are the spliceosome affinities to the canonical, alternative and cryptic splice sites, respectively; and 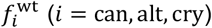 are the corresponding splice-site usage which can be read out directly from data. The affinities can then be derived from the above equation,

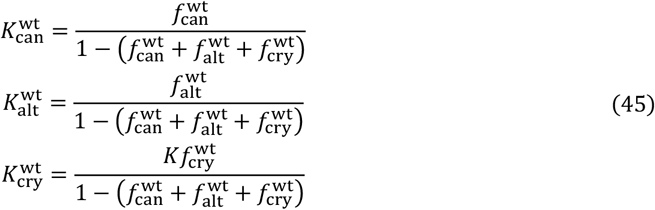

To model the effects of a hexamer motif, we assumed that the hexamer independently modulates the splice-site affinities to different extents, i.e., three new *K*s as independent free parameters. For a single hexamer motif *i* we have then,

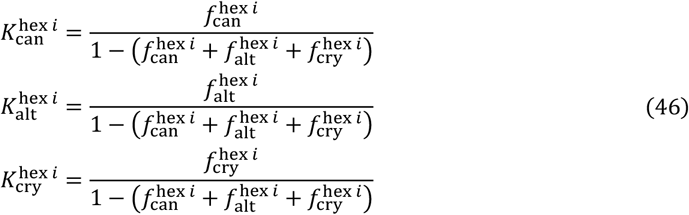

where 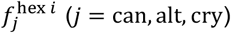 are the corresponding isoform frequences as direct readouts from data. The hexamer effect was calculated independently for the two 25-nt degenerate regions. The difference (e.g., fold change) between the inferred hexamer regulated and ‘WT’ spliceosome affinities quantifies the effects of single hexamers on splice-site choice (Fig. 5).

### Experimental *CD19* splicing analysis

#### Cell lines

HEK293 cells were obtained from DSMZ and were cultured in Gibco Dulbecco’s Modified Eagle Medium (DMEM, Thermo Fisher Scientific) with L-Glutamine + 10% Gibco fetal bovine serum (FBS, Thermo Fisher Scientific) without antibiotics. All cells were kept at 37 °C in a humidified incubator containing 5% CO_2_ and were routinely tested for mycoplasma infection.

#### *CD19* minigenes & targeted mutagenesis

In order to validate model predictions and to gain more insight in specifics of *CD19* splicing regulation, the Q5 Site-Directed Mutagenesis Kit (New England Biolabs) was used to introduce about 40 single and 16 double mutations in a *CD19* minigene, following the manufacturer’s protocol. We verified each minigene identity by Sanger sequencing (Eurofins Genomics Mix2Seq Service, following the company’s protocol). These mutations were chosen based on their model-predicted effect on (Exon2) inclusion, alt-exon2 and alt-exon3 (see Supplementary table 1 and 2 for a list of all mutations). The *CD19* minigenes consisted of *CD19* exons 1, 2 and 3 including the interspersed introns, cloned into the pcDNA3.1 plasmid vector, as previously described in Cortés-López et al. 2022^13^. The native *CD19* sequence would naturally yield almost 100% inclusion. To achieve a broader baseline of isoforms in HEK293 cells to render splicing patterns more sensitive to mutation effects, the mutations G742C and G748T were introduced around the 5’ end of intron 2, creating an artificial wildtype plasmid (referred to as “WT” in the respective figures).

#### Transfection of cells with minigenes

7×10^5^ HEK293 cells per well were seeded in 6-well TC plates prior to transfection. After 24h, minigene transfection was prepared using 2 µg of plasmid and 20 µg linear polyethyleneimine (PEI) MW 2500 (Polysciences), adding Gibco Opti-MEM (Thermo Fisher Scientific) to 100 μl total volume. After 15 min of incubation at room temperature, the transfection mixture was added dropwise to 1.9 ml fresh DMEM + 10 % FBS without antibiotics covering the cells, followed by incubation for 24 h. Only one minigene bearing a distinct single or double mutation was transfected per well, each minigene was transfected in at least n = 3 replicates.

#### Cell harvesting, RNA isolation, cDNA synthesis and RT-PCR

After 24h, cells were harvested by removing excess medium and adding 500 µL of Dulbecco’s phosphate buffered saline (DPBS, Gibco Thermo Fisher Scientific). Cells were detached from the well bottom by flushing with DPBS multiple times and transferring the cell suspension into 1.5 mL Eppendorf reaction tubes. Cells were pelleted by centrifuging at 500 x g for 5min. The supernatant was removed, and RNA was subsequently isolated using the Qiagen RNeasy Plus kit according to the manufacturer’s protocol. We modified one step of this protocol by adding only 100 to 150 µL of cell lysate to the gDNA removal columns to avoid oversaturation with genomic or residual plasmid DNA. 1000 ng of RNA were used for cDNA synthesis per sample, using the ThermoScientific RevertAid cDNA kit. We changed the second temperature step of the manufacturer’s synthesis protocol from 4 °C to 25 °C (5 min) to reduce RNA dimerization or formation of secondary structures after initial denaturation. Subsequently, 1 μl of cDNA was used as template for the RT-PCR reaction with OneTaq DNA Polymerase (New England Biolabs) at the following conditions: 94 °C for 30 s, 28 cycles of [94 °C for 20 s, 52 °C for 30 s, 68 °C for 30 s] and final extension at 68 °C for 5 min. The primers 5′-ACCTCCTCGCCTCCTCTTCTTC-3′ and 5′-GCAACTAGAAGGCACAGTCG-3′ were specific to the upstream half of *CD19* exon1 and to the downstream end of exon3, which ensured specific amplification of the five main/”canonical” *CD19* splicing isoforms (exon2) inclusion & skipping, intron2-retention and the two inclusion isoform variants alt-exon2 and alt-exon3. In indicated cases (Fig. 2f and Supplementary Fig. 5d), two additional cryptic isoforms were detected using this primer pair as well.

#### Quantification of splicing products

The *CD19* splicing isoforms amplified in the RT-PCR were identified based on their unique size using the Agilent TapeStation 4200 capillary electrophoresis system with the High Sensitivity D1000 reagents and tapes (Agilent) according to the manufacturer’s protocol. Using the peak molarity data of the TapeStation analysis software (Agilent), isoform frequencies were determined by normalizing the amount of each individual identified isoform per sample to the sum of all isoforms per sample. Per minigene, the mean of at least n = 3 samples was calculated.

Sanger sequencing (Eurofins Genomics Mix2Seq Service, following the company’s protocol) was conducted once to verify the isoform identity of quantified PCR products.

## Supplementary Figures

**Supplementary Fig. 1.**
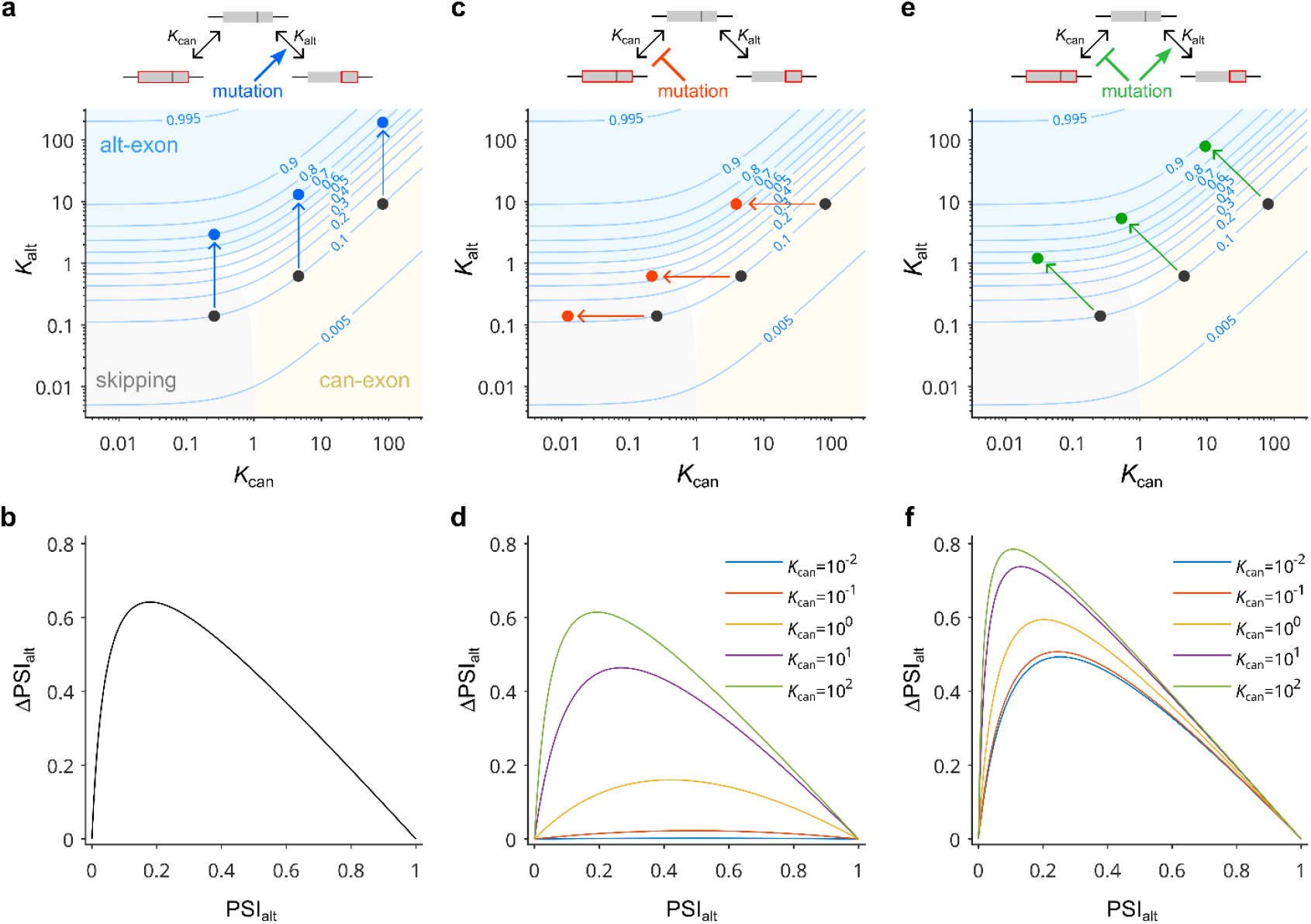
Scaling of mutation effects on splicing outcomes in multiple dimensions (top) and upon projection to the PSI space (bottom). **a** Mutation effects in the two-dimensional exon recognition space with three splice outcomes: Starting alt-exon isoform determines final alt frequency for mutations that enhance the recognition of the alt exon variant (increasing *K*_alt_). Upon this type of mutation, different starting points (black dots) in the 2D landscape lead to the same final alt isoform frequency (blue dots on the same isoline) if the starting points share the same initial alt frequency (e.g., isoline 0.1 corresponding to a frequency of 10%). Note that the starting point, however, determines the relative amount and mutation-induced shifts of canonical inclusion and skipping. **b** Effects of mutations affecting *K*_alt_ in the one-dimensional PSI projection: A single non-monotonic scaling curve governs the effects of mutations increasing *K*_alt_ (in terms of ΔPSI_alt_ = PSI_alt, mutant_ – PSI_alt, WT_ where PSI = canonical / (canonical + alternative)). Same simulations as in **a. c – f** Mutations affecting *K*_can_ (**c** and **d**) or simultaneously affecting *K*_alt_ and *K*_can_ (**e** and **f**) lead to complex scaling in the one- and two-dimensional projections. **c** Splice isoform regulation by mutations weakening the competing canonical splice site (*K*_can_). Even starting from the same alt isoform level (black dots), distinct positions in the 2D landscape, result in drastically different alt isoform frequency outcomes (red dots) for a given mutation strength (length of green arrow). **d** Scaling effects in the one-dimensional PSI projection for mutations affecting the canonical splice site (*K*_can_). Same simulations as in **c**. **e** Splice isoform regulation by mutations affecting *K*_alt_ and *K*_can_. Even starting from the same alt isoform level (black dots), distinct positions in the 2D landscape, result in drastically different alt isoform frequency outcomes (red dots) for a given mutation strength (length of green arrow). **f** Complex scaling effects in the one-dimensional PSI projection for mutations affecting the affecting *K*_alt_ and *K*_can_. Same simulations as in **e**. In a, c and e, blue isolines represent the alt-exon isoform frequency (labels). Shades indicate the regions in the 2D landscape where alt-exon (blue), canonical (yellow) and skipping (grey) isoforms are highly abundant (frequencies exceeding 50%), respectively.

**Supplementary Fig. 2.**
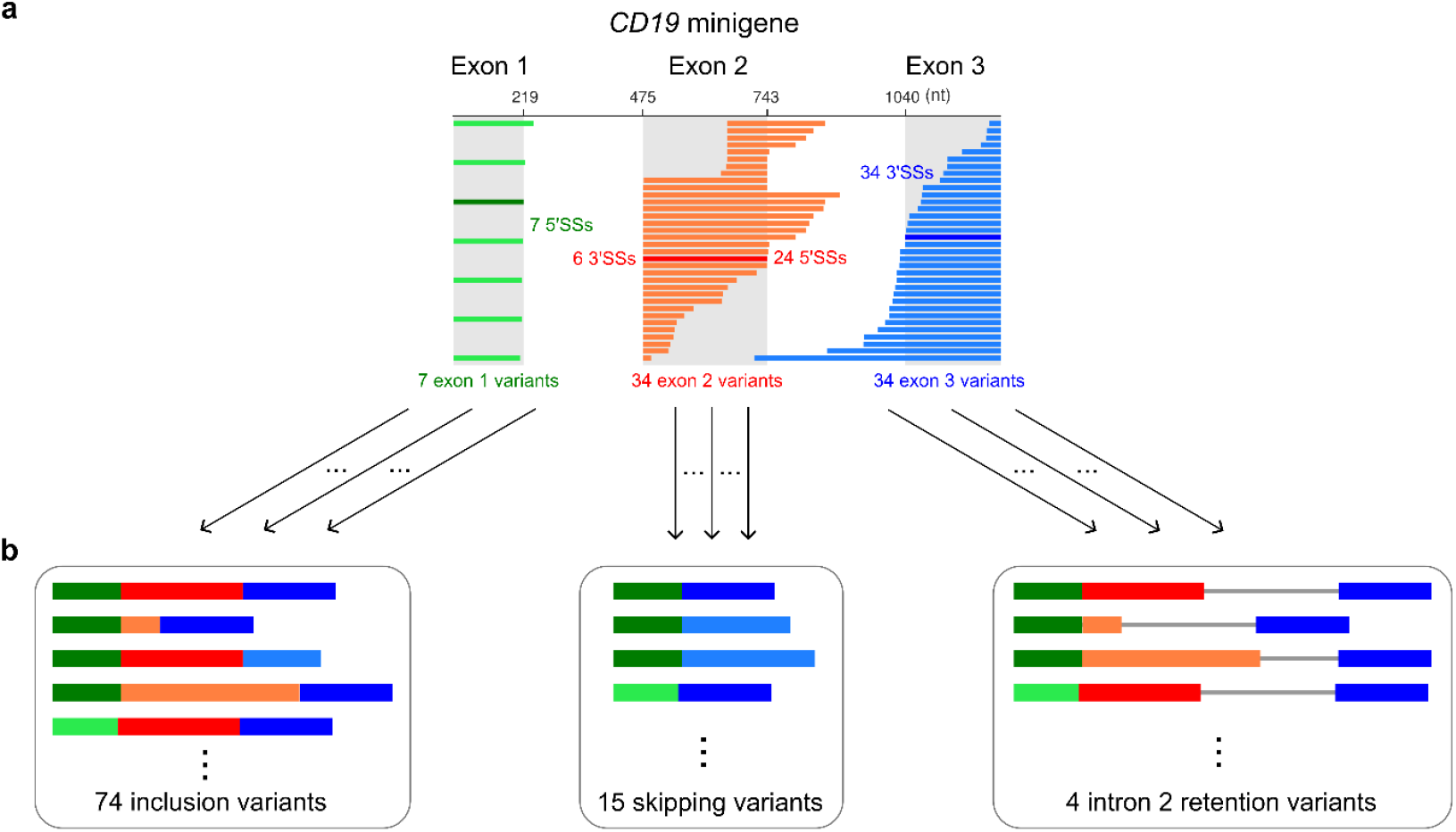
Mutagenesis data of *CD19* minigene show extensive use of alternative/cryptic splice sites. **a** A graphical summary of exon variants observed in *CD19* minigene mutagenesis data. Exon 1 contains 7 5’SSs (1 canonical and 6 cryptic sites), resulting in 7 exon variants. Exon 2 has 6 3’SSs (1 canonical, 1 alternative and 4 cryptic) and 24 5’SSs (1 canonical and 23 cryptic), which jointly form 34 exon variants. 34 3’SSs were found in exon 3 (1 canonical, 1 alternative and 32 cryptic), which leads to 34 exon variants. **b** 93 splice isoforms were studied here and they can be categorized as variants of the exon 2 inclusion (74 isoforms), skipping (15 isoforms) and intron 2 retention (4 isoforms).

**Supplementary Fig. 3.**
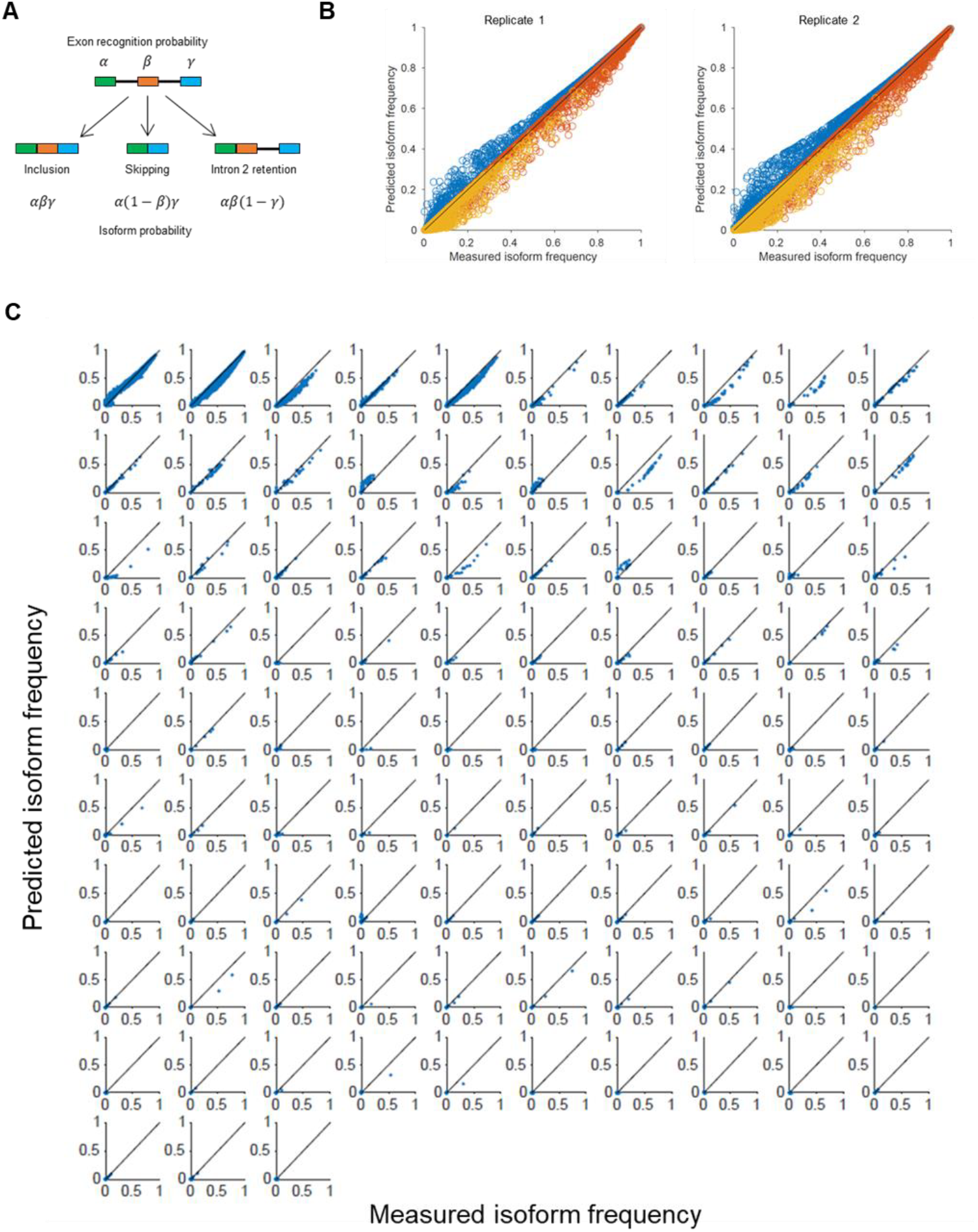
Fitting high-dimensional splice isoform measurements using maximal likelihood estimation. **a** A simplified model considering three major isoform types (inclusion, skipping and intron 2 retention) in the *CD19* minigene mutagenesis. Following exon definition model, the exon definition probabilities *α, β* and *γ* were assigned to exon 1, 2 and 3, respectively to calculate splice isoform frequencies and to construct the likelihood function for model inference of exon definition probabilities from experimentally measured isoforms frequencies (see Methods). **b** Maximum likelihood modelling quantitatively describes abundance the pooled data, in which individual isoforms were summed into three major species: inclusion (blue), skipping (red) and intron 2 retention (yellow). Shown is a comparison of model fit (y) and data (x) for each minigene in the library (dots) for two replicate experiments. **c** Extended high-dimensional model incorporating competitive alternative splice-site usage in each exon successfully reproduces the abundance of all 93 isoforms. Panels show a comparison of model fit (y) and data (x) for each minigene in the library (dots), separately for the splice isoforms.

**Supplementary Fig. 4.**
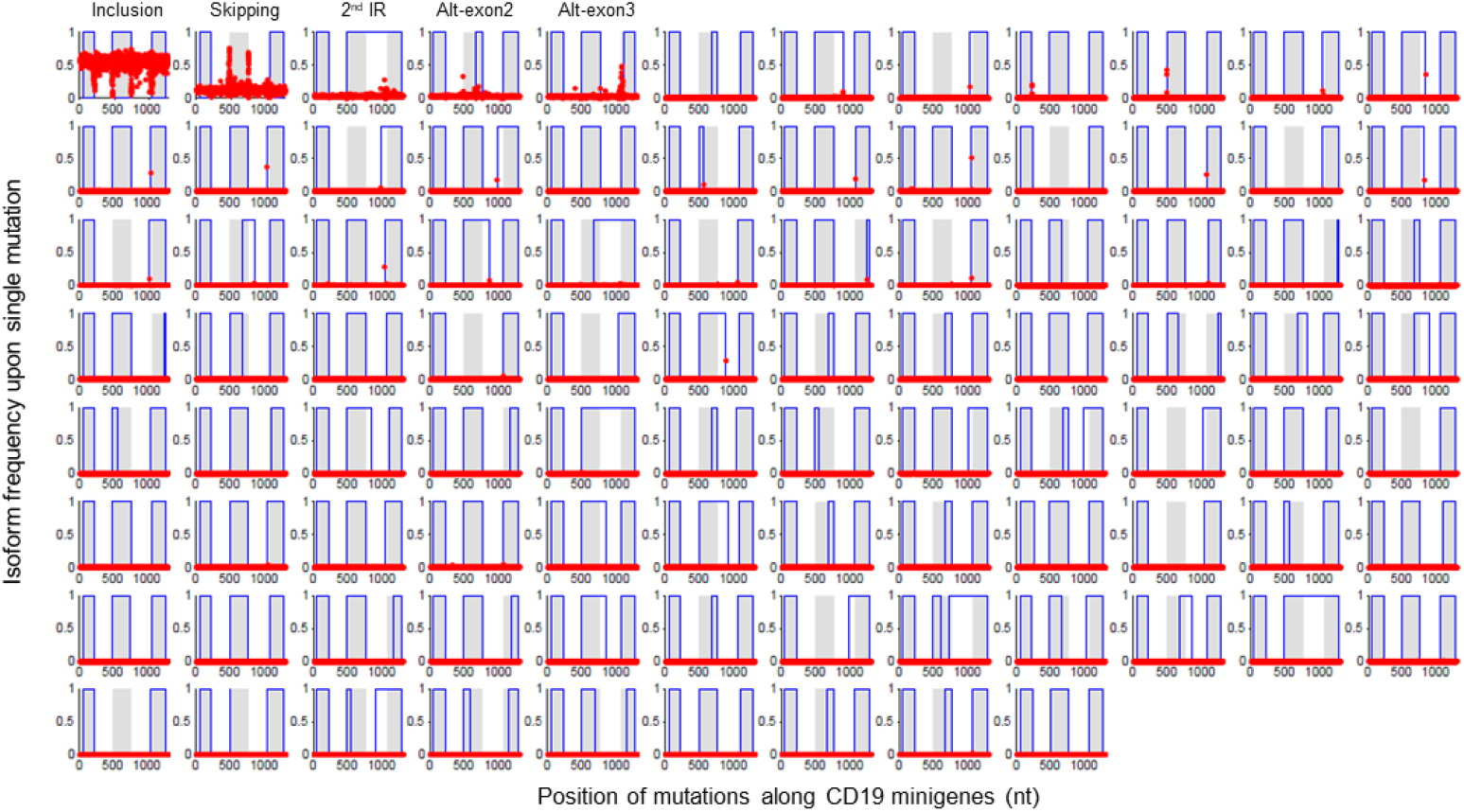
Linear regression quantifies single mutation effects on the levels of all *CD19* isoforms. Inferred splice isoform frequencies upon single point mutations (red dots) plotted against the position of the mutations in the *CD19* minigene. For reference, the grey shades represent the positioning of exon 1-3. Blue boxes indicate the splice junctions used in each isoform.

**Supplementary Fig. 5.**
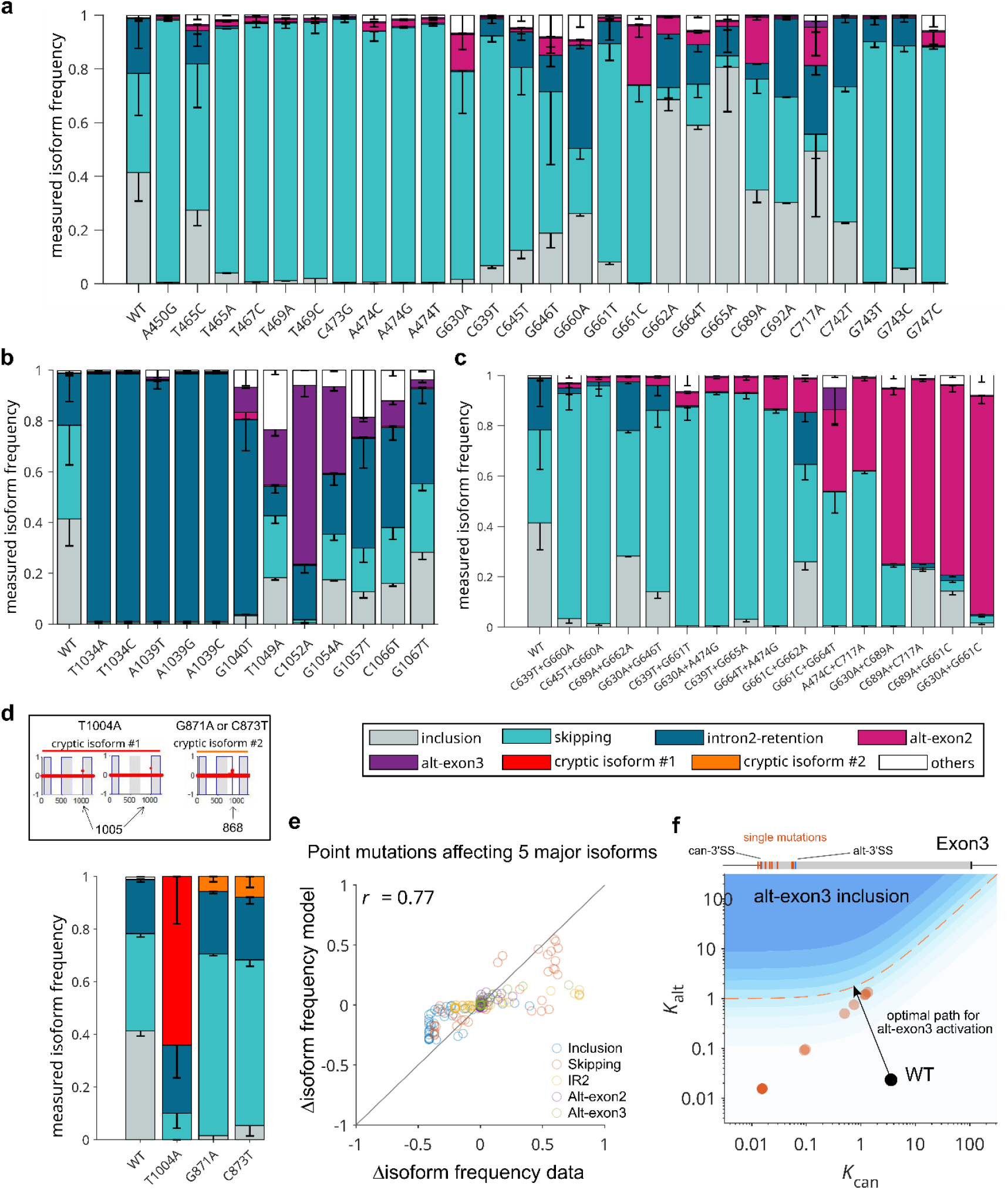
Targeted *CD19* minigene mutagenesis and splicing measurements in HEK293 cells. **a - c** Frequency of major *CD19* isoforms upon single mutations in/near exon 2 (**a**), single mutations in/near exon 3 (**b**) and double mutations in/near exon 2 (**c**). Mutations indicated at the bottom were introduced into the *CD19* minigene wildtype (“WT”, i.e., endogenous *CD19* sequence containing mutations C742T and G748C) via targeted mutagenesis. HEK293 cells were transfected with minigenes and after 24h, splicing outcomes were determined using capillary electrophoresis (see Methods). The five main isoforms and “others” (= unidentified cryptic isoforms) were quantified and expressed as isoform frequency, by normalizing individual isoform measurements to the sum of all isoforms per replicate (see legend at the bottom of the figure). All bars show means of at least n = 3 (n = 48 for the WT) with negative standard deviation as error bars. For double mutations, bars are ordered according to the isoform frequency of alt-exon2. **d** Testing model-predicted cryptic isoform activation by single point mutations. Single mutation minigenes harboring T1004A, G871A and C873T were generated using targeted mutagenesis. According to the model, these three mutations are able to create cryptic isoforms by creating/activating a novel splice site at position 1005 and 868, respectively (arrows). Mutation T1004A lengthens exon3 at its 3’end and creates two isoforms, one including and one excluding exon2, which were collectively summed up and termed “cryptic isoform #1”. Both G871A and C873T are predicted to create an isoform in which exon 2 is lengthened at its 5’end using the cryptic splice site at position 868, labeled cryptic isoform #2. The effects of these mutations on splicing were verified experimentally using capillary electrophoresis (bar graphs on the right) and data agreed well with the model prediction as shown in Fig. 2f. **e** Model-inferred mutation effects on the 5 major isoforms agree well with the validation data in HEK293 cells (same data as in **a**-**b**). **f** Targeted single mutations in exon 3 (data in **b**) were mapped in the 2D splice-site competition landscape (red dots). Black dot: WT. Black arrow: optimal path to activate alt-exon3 isoform.

**Supplementary Fig. 6.**
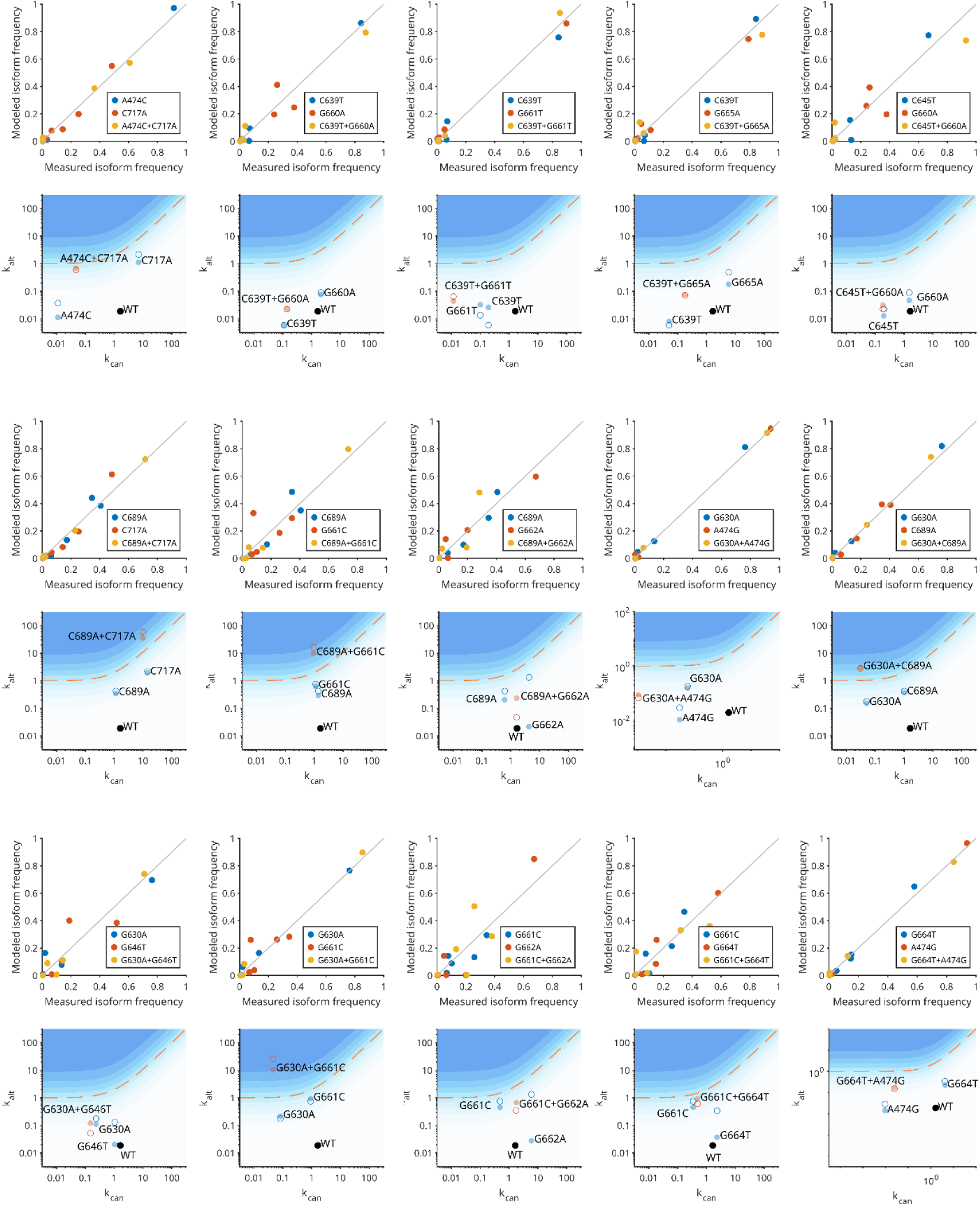
Additivity of single mutation effects quantitatively accounts for splicing outcomes for double mutations. To test how two individual mutations interact in modulating the *CD19* splicing outcome, a reduced variant of the high-dimensional splicing model was derived which describes only the 5 major *CD19* isoforms (see Methods). This reduced model was simultaneously fitted to double mutant measurements and corresponding single mutation data in HEK293 cells, assuming log-additive effects of mutations at the level of exon definition parameters (Methods). The scatter plots of 5 isoform frequencies (inclusion, skipping, IR2, alt-exon2 and alt-exon3) show the good agreement between the model (y) and data (x) for the two mutations and the corresponding combination in each panel. The inferred splice-site strengths of the canonical and alt/cry sites in exon 2 were presented in the 2D landscape. Black, blue and red circles represent WT, single and double mutations, respectively. Filled circles: inferred values by model fitting; open circles: directly derived from data. In most cases, the logarithmic 2D-landscapes show that the vector addition of single mutation effects approximately yields the double mutation effect. The merged model-data comparison for all double mutants and specific examples of 2D landscapes are shown in Figs. 3b and c, respectively.

**Supplementary Fig. 7.**
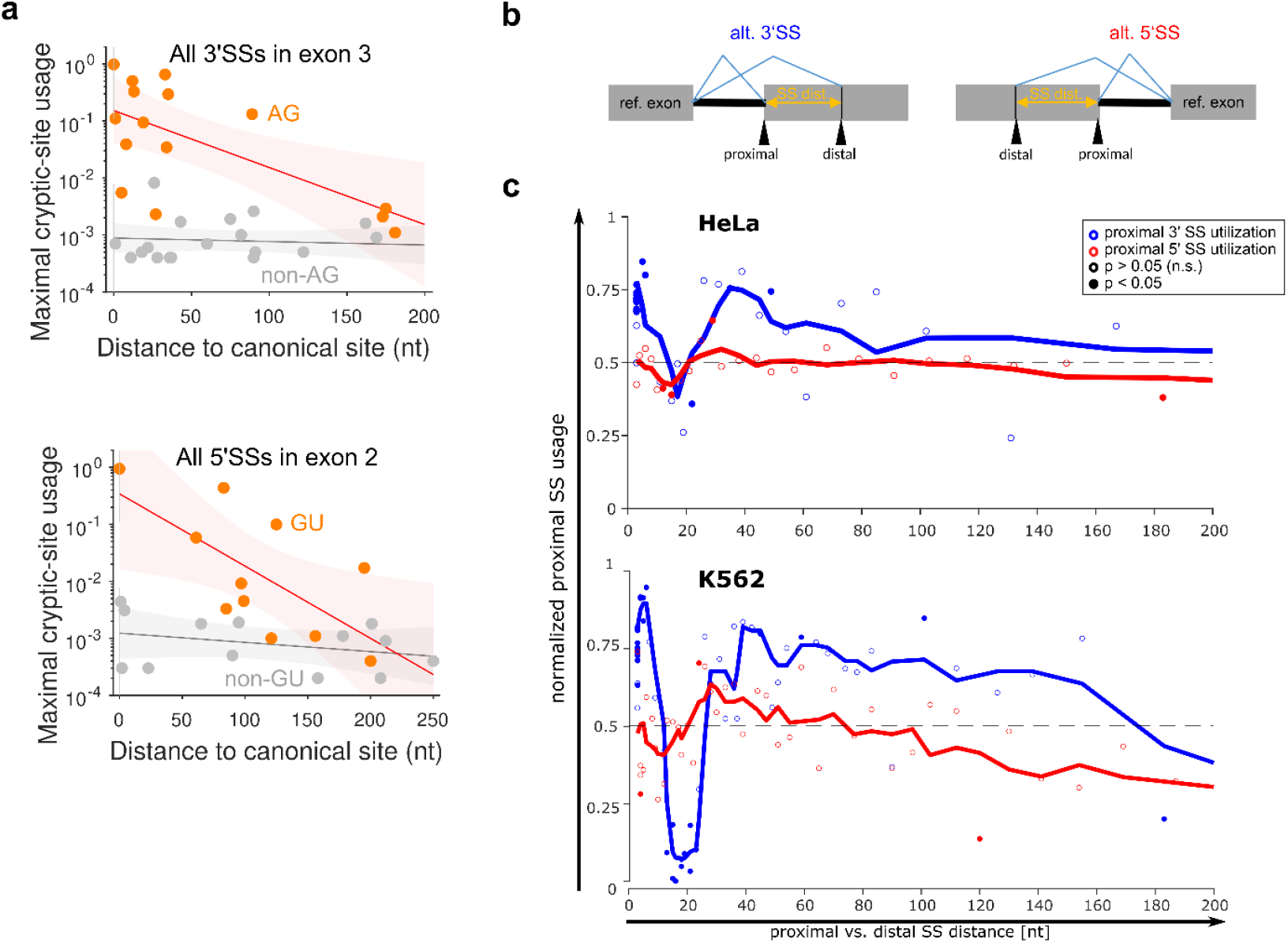
Distance dependence of alternative splice-site usage. **a** For the *CD19* mutagenesis dataset in NALM6 cells, alternative/cryptic 3’SSs (top) and 5’SSs (bottom) with consensus sequence motifs (orange dots) showed a higher maximal usage across all single point mutations in the screen when they are in close proximity to the canonical sites. The cryptic sites without consensus motifs (grey dots) are generally weakly activated across the whole *CD19* minigene mutation spectrum, independent of their positioning. **b** Analysis of distance-dependent splice site preference in genome-wide data using MAJIQ^31,32^. MAJIQ quantifies splicing outcomes in genome-wide RNA sequencing data using local splicing variants (LSVs), in which all splice junctions originating from a reference exon are collected and each junction quantified as a frequency (percent spliced-in; PSI) relative to all other events in the LSV. In the sketched examples, alternative 3’ or 5’ events are shown, for which MAJIQ designates the splice site closer to the reference exon as “proximal” and the one further away as “distal”. The distance between the two was determined and related to the utilization of either splice site. **c** Same data and processing as in Fig. 4g but without prior filtering of the data for splice site pairs with similar MaxEnt-scores.

